# Histology and spatial transcriptomic integration revealed infiltration zone with specific cell composition as a prognostic hotspot in glioblastoma

**DOI:** 10.1101/2025.10.08.681087

**Authors:** Lucas Fidon, Mélanie Lubrano, Caroline Hoffmann, Esther Baena, Céline Thiriez, Luis Cano Ayestas, Alex Cornish, Elo Madissoon, Valérie Ducret, Christian Esposito, Auriane Riou, Eric Durand, Benoît Schmauch, Elodie Pronier, Alberto Romagnoni, Almudena Espin Perez, MOSAIC consortium, Markus Eckstein, Raphaël Gottardo, Theoni Maragkou, Mohamed Shelan, Almoatazbellah Youssef, Spencer S Watson, Marc Sanson, Franck Bielle

## Abstract

**Background:** Glioblastoma (GBM), the most aggressive primary brain tumor, has a median survival of approximately 15 months. Twenty percent of patients survive beyond three years, but known clinical factors like age, performance status, resection extent, and MGMT promoter methylation status do not fully explain the observed outcomes.

**Objective:** Our objective was to identify novel histology derived biomarkers associated with end-of-spectrum overall survival (OS) to provide novel biological insight with a translational potential.

**Methods:** We analyzed a total of 748 GBM patients from 3 different cohorts, uniquely enriched in long survivors (n=98 with overall survival (OS) > 5y including n=196 with OS≥3y), with clinical data and H&E slides obtained from the primary tumor at baseline. We propose an interpretable machine learning (ML) methodology for the discovery of histological biomarkers. Our method learned to segment each H&E slide into three distinct regions associated with long-term survival, short-term survival, and non-informative tissue. We characterized these regions by integrating unsupervised learning, nuclei segmentation, blood vessels detection, pathologist annotations, and multimodal data including spatial transcriptomics from n=31 patients of the GBM MOSAIC dataset to discover fully interpretable biomarkers.

**Results:** Our OS prediction model using histology and clinical data as input achieved an area under the curve (AUC) of 0.85 for the classification of patients between OS<2 and OS≥3y in external cohort validation, outperforming significantly models trained on clinical data or on histology alone (AUC of 0.76; 0.73, respectively).

Two novel biomarkers were predicting poor survival: the presence of regions of lowly infiltrated white matter enriched in malignant cells with a mesenchymal-like phenotype, and lower levels of angiogenesis associated with higher hypoxia response in the main tumor regions. We also found that a subtype of immunosuppressive tumor macrophages - defined by high *PLIN2* expression and lipid accumulation- is consistently enriched in histological areas predictive of poor prognosis.

**Conclusion:** Our interpretable ML methodology identified a novel prognostic impact of biological processes and cell types according to distinct tumor regions of GBM. These results pave the way for spatially-informed biomarkers to improve risk stratification and for personalized spatially-targeted therapeutic strategies.

**Key highlights:**

1. Our ML model identified histological biomarkers predicting prognosis independently from known clinical factors
2. The region of lowly infiltrated white matter enriched in malignant cells including a mesenchymal-like phenotype is predictive of poor prognosis
3. Angiogenesis is increased in areas predictive of long survival in main non-necrotic tumor regions.
4. The subtype of macrophages expressing *PLIN2* and associated with increased lipid metabolism was associated with poor prognosis in all GBM regions.

**Highlights:** 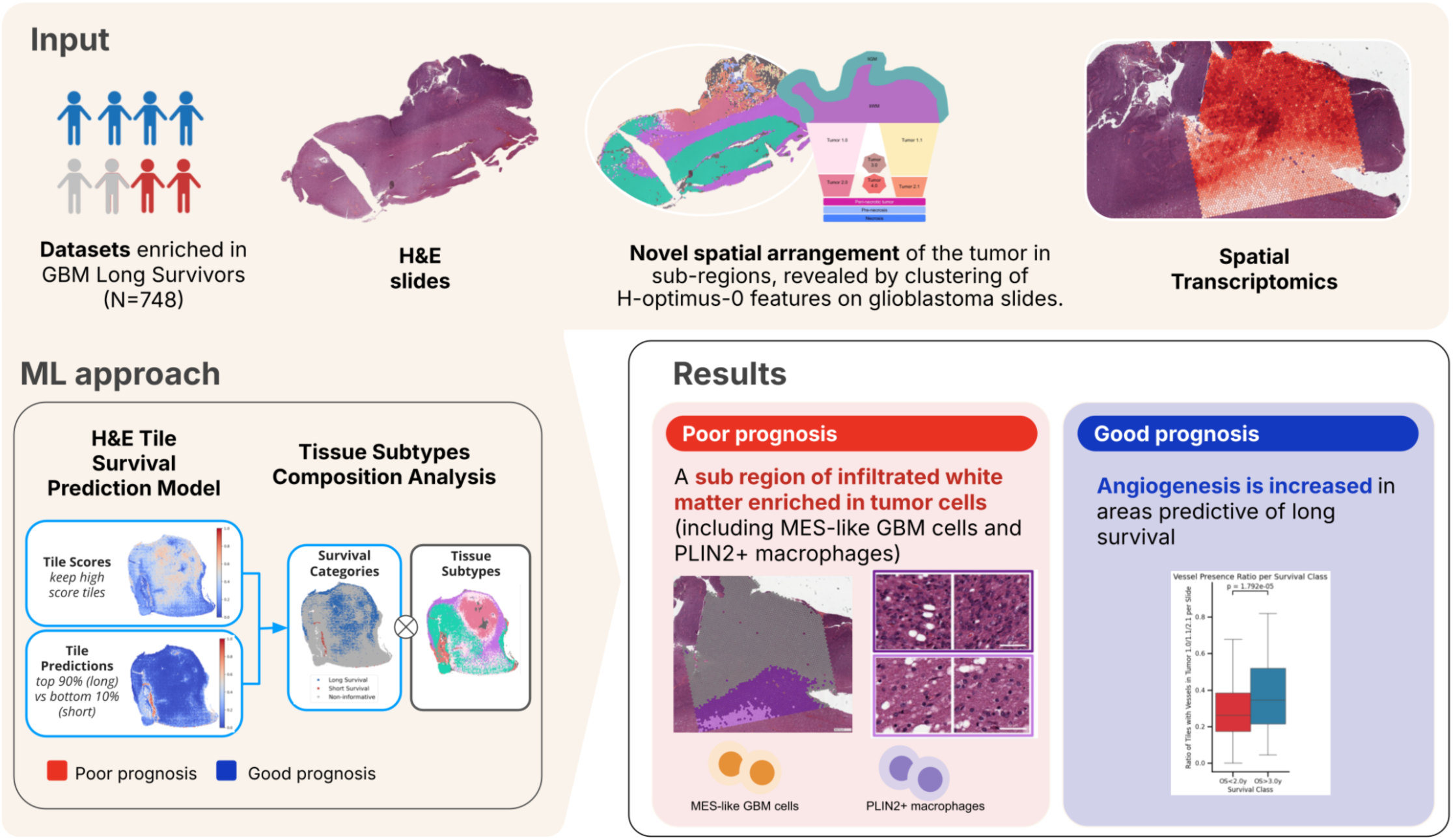

## Introduction

Glioblastoma (GBM) is the most common and aggressive primary brain tumor in adults. It corresponds to the histomolecular diagnosis of glioblastoma, IDH-wildtype grade 4 according to the reference WHO 2021 classification. It is characterized by rapid progression and a median overall survival (OS) of approximately 15 months and only around 20% of patients are classified as long survivors (survive more than 3 years) when treated with the standard of care (SOC), including maximal safe resection and concomitant radiochemotherapy with temozolomide (TMZ) (Ostrom et al. 2022; Stupp et al. 2009; Chinot et al. 2014; Thomas-Joulié et al. 2024; Gilbert et al. 2013). Long range infiltration of brain parenchyma by isolated tumor cells makes it particularly challenging to surgically resect the tumor and to target the residual disease by adjuvant treatment, which remains a major barrier to tumor eradication and achieving a definitive cure for GBM.

Adverse clinical prognostic factors are male gender, greater age, lower postoperative Karnofsky Performance Status (KPS), and larger residual tumor volume (Karschnia et al. 2025). O6-methylguanine-DNA methyltransferase promoter methylation status (MGMT status) remains the only biomarker currently used to guide therapeutic decisions and predicts better prognosis and better response to TMZ (Hegi et al. 2005). However, the absence of a standardized definition and assay to assess MGMT promoter methylation status limits its ability to distinguish between poor and good responders.

Histopathology of GBM is characterized by poorly differentiated tumor cells of a predominant astrocytic phenotype encompassing a wide range of morphological variants with intratumoral and interindividual heterogeneity. Some histolopathological variants of GBM are associated with distinct oncogenic drivers: the “small cell glioblastoma” variant is typically associated with *EGFR* amplification present in 40% of GBM while distinct features (regular ovoid nuclei, vague pseudorosetting, dense and branched capillary network, microcalcifications) are associated with *FGFR3–TACC3* fusion present in 3% of GBM and exclusive with *EGFR* amplification (Louis et al. 2007; Di Stefano et al. 2020; Bielle et al. 2018). While high mitotic activity, microvascular proliferation (MVP), and palisading necrosis are hallmarks of GBM used for diagnosis, they are not predictive of OS (Gilard et al. 2021; G. Linkous et M. Yazlovitskaya 2011). Identifying histological markers associated with OS could suggest new stratifications of patients in future clinical trials and refine our understanding of GBM mechanisms of resistance to SOC treatment (Hegi et al. 2005; Gorlia et al. 2008).

The use of machine learning (ML), and in particular multiple instance learning (MIL), to predict survival using digitized H&E of primary GBM cases acquired at the time of diagnosis has already been proposed in the literature (Mobadersany et al. 2018; Di et al. 2022; Carmichael et al. 2022; Redlich et al. 2024; Verma et al. 2024; Luo et al. 2023; Yin et al. 2024; Baheti et al. 2024; 2025; Saada et al. 2024). Promising results were obtained in terms of survival prediction performance as measured by the C-index metric. However, those studies were primarily interested in benchmarking deep learning methodological novelties and did not explore the discovery of novel histological biomarkers (Redlich et al. 2024). Our work extends beyond such approaches by developing an interpretable ML method specifically designed to identify new imaging biomarkers and associate them with underlying tumor biology.

Here, we leveraged a novel dataset of 748 GBM patients with a specific enrichment in long survivors. We developed an interpretable histology-based model predicting end-of-spectrum survival to discover which histological features differed between short (OS<2y) and long (OS≥3y) survivors. To gain biological insights, we defined a novel unsupervised classification of histological regions in GBM, and identified 10 relevant regions further annotated by neuropathologists. We also leveraged spatial transcriptomics aligned with their histology slides from 31 patients from the GBM MOSAIC consortium dataset (MOSAIC Consortium et Hoffmann 2025) to determine the cell subtypes and pathways enriched in the areas predicted as associated with short or long survival. We identified two novel histology-derived biomarkers predicting poor survival in GBM: (i) presence of areas within the lowly infiltrated white matter enriched in malignant cells with a mesenchymal-like state; and (ii) lower levels of angiogenesis, endothelial cells and pericytes within non-necrotic tumor regions. Additionally, a subtype of macrophages expressing *PLIN2* was increased in histological areas predictive of poor prognosis throughout most GBM regions.

## Results

### Histological features are complementary to routine variables to predict end-of-spectrum survivals

#### A glioblastoma cohort enriched in long-term survivors and reflecting known prognostic factors

A novel dataset with 802 patients from 3 centers (center A (Hôpital Universitaire de la Pitié Salpêtrière AP-HP, Paris, FR), center B (Universitätsklinikum Erlangen, DE), center C (Insel Gruppe AG, Bern, CH)) was collected along with clinical information including outcomes and histological slides of the tumor before any treatment (<u>Methods</u>). Following slide review and quality controls, 748 patients remained eligible for inclusion (Fig 1A, Suppl. Table A). This dataset is uniquely enriched in long survivors (196 patients with OS > 3 y (years), including 98 patients with OS > 5 y), representing 24 to 48% of the population in the 3 centers, whereas the expected percentage is only around 20% (Stupp et al. 2009; Chinot et al. 2014; Thomas-Joulié et al. 2024; Gilbert et al. 2013).

**Figure 1.**
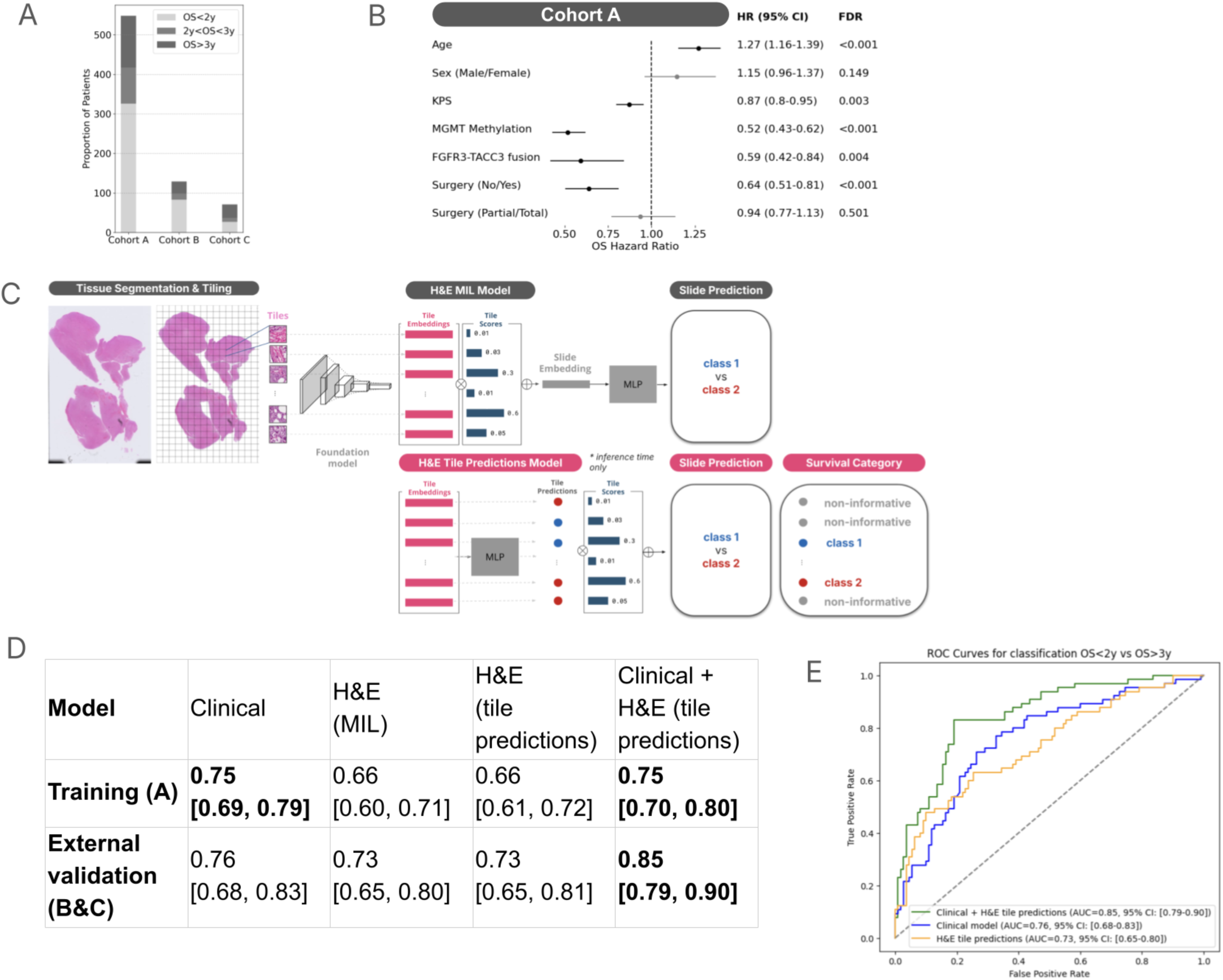
Overview of the training and testing cohorts and performance of the model showing histology and clinical features are complementary to predict end-of-spectrum survivals. **(a)** Number of patients with GBM in all cohorts with H&E slides meeting the quality control assessment (outlined in the Methods section) and partition into short (OS<2y), medium (2y<OS<3y) and long survivors (OS≥3y). **(b)** Univariate analysis: hazard ratio for OS on the cohort A for clinical parameters known to be associated with OS in the SOC for GBM (n = 548 patients). **(c)** Illustration of the H&E MIL model and H&E tile predictions model. **(d)** ROC AUC (mean ± 95% CI via bootstrapping) for the clinical model (logistic regression ensemble using MGMT, age, surgery status, KPS, and gender), the H&E MIL model, the H&E tile prediction model, and their combination. **(e)** ROC curve of our H&E tile predictions model, clinical model and the combination of H&E tile predictions model and clinical model on the combined testing cohorts. KPS: Karnofsky Performance Status, MGMT: O6-methylguanine-DNA methyltransferase promoter.

First, we performed an univariate analysis of clinical and molecular parameters in the full cohort A (n = 548 patients) and in the external validation sets (cohort B&C; n = 200 patients). Cohorts B and C were combined to increase statistical power. The median age was 60.5 (52.4-67.2) in cohort A and 61.6 (53.9-70.5) in cohort B&C. The sex ratio (M:F) was 1.51 (329:218) in cohort A and 1.25 (111:89) in cohort B&C. The respective proportions of resection and biopsy were 81.8% and 16.8% in cohort A and 100% and 0% in cohort B&C. The respective percentage of methylated versus unmethylated MGMT status were 50.5% and 46.2% in cohort A and 55.5% and 44.0% in cohort B&C. Some clinical data were missing (Suppl. Table B). The training and the external validation sets were mostly comparable, with comparable age range and MGMT methylation status proportions. The main clinical difference was that 17% of the patients in the training set did not undergo surgery, as opposed to none in the external validation set. Karnofsky Performance Status (KPS) and FGFR3-TACC3 fusion status were missing in the external validation set. Five clinical and molecular parameters were found to be significantly associated with better OS in univariate analysis in cohort A: lower age at diagnosis, higher KPS, positive MGMT methylation status, presence of a FGFR3-TACC3 fusion, and resection versus no resection (Fig 1B). Similar findings were obtained in the external validation set for the variables present (Suppl. Fig 1A). Male sex was not a prognostic factor in these two datasets. There was no significant correlation between these clinical and molecular parameters (Suppl. Fig 1B). Those findings were consistent with known prognostic biomarkers (Hegi et al. 2005; Lacroix et al. 2001; Liu et Wang 2025).

#### Interpretable AI-based methodology to localize histology regions associated with survival

We developed a prognosis model based only on histological features. We designed a prediction task for end-of-spectrum survival classification, predicting the binary labels of OS < 2y (thereafter named ‘short survivors’) versus OS ≥ 3y (‘long survivors’). Patients with intermediate survival (2y≥OS>3y, n=121) were removed to maximise the biological differences. We included 632 patients: 457 from center A for model training, and 175 from centers B and C for external validation. In addition, here to confirm the methodology used for the model, and thereafter to confirm the validity of our interpretability method in brain tissue, we developed a second model to predict the binary label of FGFR3-TACC3 fusion status: positive versus negative. For this task, we included 440 patients from center A (44 fusion-positive and 396 fusion-negative) with model training performed using cross-validation (Suppl. Table C, <u>Methods</u>).

Our method was based on deep learning and multiple instance learning (MIL) with four main steps (Fig. 1C). Firstly, the whole slide images (WSI) were tessellated into tiles of size 224x224 pixels at a resolution of 0.5 micron per pixel and each tile transformed into an embedding vector of 1536 features using the histology foundation model H-optimus-0 (Saillard et al. 2024). Secondly, tiles with non-tumoral content were filtered out using an automatic tissue segmentation method to reduce noise and improve performance (Suppl. Table D, <u>Methods</u>). Thirdly, an ‘H&E MIL’ model based on a deep learning architecture with an attention mechanism was trained using the patient-level binary label (<u>Methods</u>). Fourthly, at inference, the order of operation of the prediction layer and the aggregation layer were inverted in the H&E MIL model. The resulting model was named the ‘H&E tile predictions’ model (<u>Methods</u>). The H&E tile predictions model is interpretable, providing two outputs for each histology tile: a *tile score* (indicating its contribution to patient-level prediction) and a *tile prediction* (indicating predicted class association). By combining these two outputs, we categorized tiles into three classes relevant for each binary classification task. For the end-of-spectrum survival prediction (‘long survival’ with OS >3y vs ‘short survival’ with <2y), tiles were categorized as associated with ‘long survival’ (high tile scores; high tile prediction); ‘short survival’ (high tile scores; low tile prediction), and non-informative tiles (all others) (<u>Methods</u>). Similarly for FGFR3-TACC3 fusion status prediction, tiles were classified as associated with the presence of the fusion (high tile scores; high tile prediction), the absence of the fusion (high tile scores; low tile prediction), and non-informative tiles (all others). This framework allowed spatial localization of biologically relevant features and provided insights into how the model generated patient-level predictions. Unlike standard H&E-based MIL models, which produce only tile-level scores without association to the classification labels, our approach incorporates predictive context, enabling biologically meaningful interpretation.

#### H&E tile predictions and H&E MIL models achieve the same performance

We evaluated the performance of both our H&E models (MIL and Tile prediction) for GBM patients, despite the shown interpretability differences. Because the H&E MIL model included non-linear MLP layers after attention-based aggregation (<u>Methods</u>), it is not mathematically equivalent to the Tile prediction model, necessitating direct performance comparison. For end-of-spectrum survival prediction, both models performed similarly in our external validation cohort: both had ROC AUC of 0.73 with 95% confidence intervals (CI) of [0.65–0.80] for H&E MIL and [0.65–0.81] for H&E Tile prediction, and DeLong test was not significant (p=0.92) (Fig. 1D, Suppl. Table E, Suppl. Table F). Likewise, for prediction of the FGFR3-TACC3 fusion status, the H&E MIL and H&E Tile predictions models also achieved nearly identical performance in cross-validation (ROC AUC of 0.89 [95% CI: 0.83–0.95] vs 0.90 [95% CI: 0.84, 0.95]; Suppl. Fig 2).

In conclusion, on both tasks, the H&E MIL model and the H&E tile predictions model achieved nearly identical performance, indicating that the inversion of operations (as described in step (iv) above) in the H&E tile predictions model did not compromise predictive accuracy. This equivalence in performance supports the downstream use of the H&E Tile Predictions model for histological interpretation.

#### Histological features are complementary to clinical variables to predict end-of-spectrum survival

To evaluate the added prognostic value of our H&E tile predictions model as compared to the routine clinical parameters, we developed a multivariate clinical model for our OS classification task based on MGMT promoter methylation status, age, KPS, surgery status (none, partial, total), and sex (Fig. 1B; <u>Methods</u>). We compared the clinical model, the H&E tile predictions model, and a multimodal model resulting from the combination of the clinical and H&E models obtained by taking the average of their predictions (Fig 1D-E; Suppl. Table E). The multimodal model achieved an AUC of 0.85 in external validation, outperforming significantly the clinical model and the H&E tile prediction model (AUC of 0.76; 0.73, respectively). Similar results were obtained with the other 2 metrics (PR AUC, and C-index) (Suppl. Table E).

Altogether, our results show that, in the population undergoing optimal standard of care (surgical resection and adjuvant therapy as recommended), the combination of a clinical model limited to four variables (MGMT promoter methylation status; age; gender; partial vs total surgery) and H&E tile prediction model provided the best prediction for survival. This demonstrates that the H&E tile model provides novel and complementary information to known clinical prognostic factors.

### AI-based identification of glioblastoma tissue regions and their association to survival

Next, we aimed to identify histological features associated with end-of-spectrum survival by using the intrinsic interpretability capacity of our H&E tile predictions model. We performed two preliminary tasks for that purpose: (i) we demonstrated the ability of our model to uncover relevant histological patterns in brain tissue on the molecular use case of FGFR3-TACC3 fusion prediction, (ii) we proposed a novel classification of GBM tissue regions and, to advance towards interpretation, we assessed the distribution of short and long survival tiles in these regions.

First, to demonstrate the validity of our interpretable model in brain tissue, we evaluated its ability to identify established histological biomarkers, using the FGFR3-TACC3 fusion, a molecular alteration known to be linked to distinct histopathological features (Bielle et al. 2018) (Suppl Fig 3A-B). To this end, 83 regions of 336μm x 336 μm were selected according to their FGFR3-TACC3 fusion-positive or fusion-negative tile predictions. An expert neuropathologist (FB), blinded for the samples’ molecular status and tile predictions, assessed each region for hallmark FGFR3-TACC3-associated morphologies (Suppl. Fig 3C; <u>Methods</u>).

As expected, a significant enrichment of known FGFR3-TACC3 histological features such as monomorphous ovoid nuclei, endocrinoid network of thin capillarities, nuclear palisading, pseudorosette and calcification were found in fusion-positive regions (Suppl. Fig 3D, <u>Methods</u>). In addition, based solely on histological evaluation of the H&E regions, the pathologist was able to correctly infer FGFR3-TACC3 fusion status in 49 regions with 100% accuracy matching both molecular ground truth and tile predictions (Suppl. Fig 3E), thereby confirming that the model identifies the same discriminative histological patterns used by expert pathologists.

Altogether, using FRGFR3-TACC3 fusion as a use case, we demonstrated the model’s capacity to identify established, biologically grounded features predictive of molecular status and interpretable histological regions.

Second, we developed a novel classification of GBM tissue regions by extracting features with the H-optimus-0 model (Fig 2A) followed by unsupervised clustering, which identified 14 distinct tissue regions (Suppl. Fig 4, <u>Methods</u>). The regions were discovered on the training dataset and then transferred to the external testing dataset, ensuring the same regions were identified in all the H&E slides across the datasets. Two neuropathologists reviewed the 14 tissue subtypes for consistency and provided nomenclature and description for each (Fig 2A; Table 1; Supp Fig 5, Supp Fig 6A, <u>Methods</u>). Moreover, to further characterise these regions, we used matched histology slides, single-nuclei transcriptomics (SnRNAseq, by 10x Chromium) and Spatial transcriptomics (SpT, by 10x Visium SD) of 31 patients with GBM from the MOSAIC consortium (MOSAIC Consortium et Hoffmann 2025) to assess the distribution of 41 cell types and states within each region (Fig 2B; <u>Methods</u>; Table H). The identified regions were found to be distinct spatially localized regions on the slides (Suppl Fig 6B, <u>Methods</u>), with several slides sharing similar spatial organization (see Fig 2C; Suppl Fig 6C). In particular, we identified a region corresponding to the cerebral cortex tissue with the highest presence of neurons and a low level of tumor infiltration that was named lowly-infiltrated grey matter (liGM), and similarly a second region of liWM. Five additional regions, representing infiltrated tumor tissue (designated Tumor 1.0, 1.1, 2.0, and 2.1,2.1 and “rare tumor phenotypes”) were characterized according to their histological specificities, tumoral cell density, and oedematous extracellular spaces. Specifically, Tumor 1.0 and 1.1 had more oedematous extracellular space than Tumor 2.0 and 2.1 regions, and Tumor 1.1 and 2.1 had higher tumor cell density respectively than Tumor 1.0 and 2.0. A region exhibiting significant MVP and endothelial hyperplasia was designated as “Perinecrotic tumor”. Artefacts, irrelevant and/or tumoral tissue with technical alteration were found to dominate four other clusters: “Hemorrhage”, “Artefact”, Tumor 3.0 and Tumor 4.0. Two regions representing distinct stages of necrosis were identified: the “Necrosis” region and the “Pre-necrosis” region, both displaying eosinophilic necrosis, with the latter also displaying nuclear debris or pyknotic nuclei. The division of the regions into tumor, necrosis, and lowly-infiltrated brain regions was further validated by blinded slide-level annotations (<u>Methods</u>). Finally, after exclusion of necrotic and artefacted regions, ten meaningful regions were retained for downstream interpretation.

**Figure 2.**
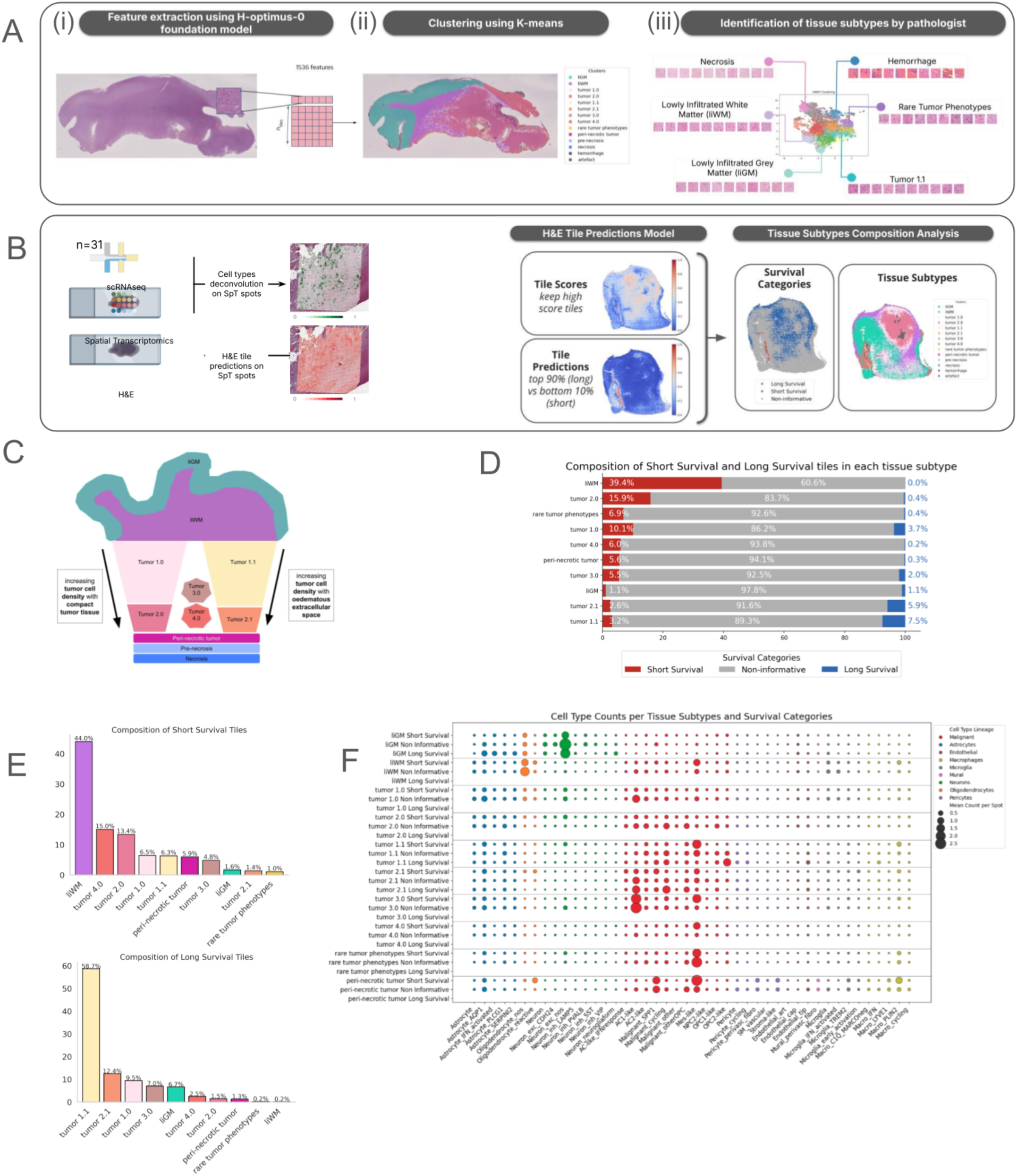
Interpretable AI models identify histological subtypes associated with poor survival. **(a)** Overview of the clustering approach and characterization of identified regions. **i)** Features are extracted from each tiles of H&E slides using the histology foundation model H-optimus-0. **ii)** Tile features from each slide are clustered using K-means, trained on the discovery cohort (cohort A) and applied to the validation cohorts (cohorts B and C). **iii)** Expert neuropathologists review and annotate each cluster to define distinct regions. **(b) Methods overview. Left**: Graphical representation of the methods used to align paired H&E and Visium spatial transcriptomics for cell type annotation in 31 samples from Cohort B from the MOSAIC consortium. **Right:** Overview of the method used to categorize tiles according to their tile score and tile prediction in 3 classes: high score + high prediction = Long Survival, high score + low prediction = Short Survival, all others = Non-informative. **(c)** Schematic representation of common spatial arrangements of the regions within tumor sections. **(d)** Proportion of tiles associated with short survival, long survival, and non-informative categories in the validation cohort in the different regions (excluding the 4 regions with necrosis, hemorrhage, and artefacts) (cohorts B and C). **(e)** Distribution of regions among Long Survival and Short Survival tiles in the validation cohorts (cohorts B and C). **(f)** Average of cell count per cell type in the different tissue regions as estimated using the output weights of the cell2loc deconvolution algorithm and 31 samples from cohort B with paired H&E, Visium spatial transcriptomics, and scRNAseq. Rare cell types (median counts < 0.1) were excluded.

**Table 1.** Histological description of the tissue regions. Identified tissue subtypes annotated by an expert neuro-pathologist (FB). Supplementary Figure 5 provides illustration for each tissue subtypes.

| Region name | Histology | Tumor cell density | Oedematous extracellular spaces |
| --- | --- | --- | --- |
| Lowly Infiltrated Grey Matter (liGM) | Cerebral cortex with or without low tumor infiltration | no / low | no / low |
| Lowly Infiltrated White Matter (liWM) | White matter with or without low tumor infiltration | no / low | no / low |
| Tumor 1.0 | Tumor tissue of moderate cellularity with few oedematous extracellular spaces | low / moderate | low |
| Tumor 1.1 | Tumor tissue of moderate cellularity with numerous oedematous extracellular spaces | moderate / high | moderate |
| Tumor 2.0 | Dense tumor tissue without oedematous extracellular spaces | moderate / high | no |
| Tumor 2.1 | Dense tumor tissue with few oedematous extracellular spaces | high | low |
| Tumor 3.0 | Tumor with smaller nuclei and few densely stained nuclei (with technical alterations) | low / moderate / high | no |
| Tumor 4.0 | Tumor with elongated/ovoid, smaller and darker nuclei (with technical alterations) | low / moderate / high | no / moderate / high |
| Perinecrotic tumor (and blood vessels) | Perinecrotic tumor enriched in neoangiogenesis and microvascular proliferation |  |  |
| Rare tumor phenotypes | Rare tumor phenotypes (eg: tumor cells with prominent nucleoli) |  |  |
| Necrosis | Eosinophilic necrosis |  |  |
| Pre-necrosis | Few nuclei within eosinophilic necrosis |  |  |
| Hemorrhage | Hemorrhage |  |  |
| Artefact | Artefact |  |  |

For each of these ten regions, we assessed the distribution of tiles associated with short survival, long survival or non-informative tiles (Fig. 2D-F; Supp Fig 7). Among all regions, liWM uniquely and consistently exhibited the lowest proportion of non-informative tiles (∼60%) and 100.0% of informative tiles were linked to short survival in both datasets (Fig. 2D-F; Supp. Fig. 7A). This made liWM the only tissue region reliably and specifically associated with poor prognosis. In contrast, Tumor 1.0, Tumor 1.1, and Tumor 2.1 were the most frequent tissue subtypes in long survival associated tiles (Fig. 2E; Supp. Fig. 7B), but their compositions were heterogeneous, containing a mixture of long and short survival tiles as well as non-informative ones. The remaining six regions had a similar composition of long and short survival tiles (Fig. 2E; Supp Fig 7). We assessed the prognostic value of those individual regions by using the proportion of liWM, Tumor 1.0, Tumor 1.1, and Tumor 2.1 per slide as candidate biomarkers (Supp Fig. 8A). However, none outperformed random prediction, likely due to the substantial presence of non-informative tiles within these regions. After exclusion of non-informative tiles, the proportion of liWM achieved statistically significant, non-random performance in predicting short survival (p < 0.01) (Supp Fig 8B). This finding motivated a refined interpretation using the comparison of informative versus non-informative tiles within each region, which is developed in the next result sections.

Finally, using the SpT data, we determined the distribution of malignant and non-malignant cell types in each of the ten relevant tissue regions and by the three tile categories (predictive of short and long survival, and non-informative) (Fig. 2F). Spatially aligned histology slides and SpT data enabled the extraction of histology tiles centered on each SpT spot. Using the H&E tile predictions model, each spot was categorized as belonging to a given region as listed in Table 1, and associated with short survival, long survival, or non-informative. We then estimated cell type composition per spot. Overall, malignant cells were the predominant cell type across most tumor regions (liWM, Tumor 1.0-4.0 and Peri-necrotic tumor), with relatively lower abundance in the liWM and Tumor 1.0 regions. The lowest average malignant cell count was observed in the liGM region. For non-maligant cells, the liGM cluster was the most neuron-enriched tissue region. Both the liGM and liWM regions were enriched in oligodendrocytes, though the average cell count was significantly higher in liWM *(p < 0.001)*. Microglia were most abundant in the liWM and Perinecrotic tumor regions. Macrophages exhibited a relatively uniform distribution across all tissue regions, with the exception of liGM, where they were notably absent. Astrocytes and endothelial cells were found in similar abundance across all tissue regions, with the exception of Tumor 4.0 and Rare tumor phenotypes. Consistent with current literature (Neftel et al. 2019; Wang et al. 2017), we observed that Mesenchymal-like (Mes-like) and *SPP1*-expressing malignant cells, were preferentially associated with short survival in the tumor regions. Mes-like cells are known to be associated with hypoxia and found in the tumor core (Neftel et al. 2019), as well as malignant cells highly expressing and secreting *SPP1* (osteopontin), key driver of the mesenchymal state and cell communication with surrounding immune cells (Pombo Antunes et al. 2021; Kijewska et al. 2016). A key finding was the identification of Mes-like and *SPP1*-malignant cells within the regions of liWM predictive of short survival. Second, we identified a subset of macrophages expressing *PLIN2* (Macro_PLIN2) which was specific to short survival in most regions (Table H).

### Identification and characterization of a new biomarker within infiltrated white matter predictive of short survival

To further understand the different biology between the tiles found associated with short-survival versus non-informative in liWM, we performed comparative analyses on histological features and SpT. As previously noted, no tiles associated with long survival were found in liWM.

First, we quantified nuclear morphology across all H&E tiles in the training cohort using a deep learning–based nuclei segmentation pipeline (see <u>Methods</u>). Two cell-level features were extracted per tile: nuclear density (number of nuclei per tile) and nuclear size (median nuclear area in µm²) (Fig. 3A). In liWM, tiles associated with short survival exhibited significantly higher nuclear density and smaller nuclear sizes compared to non-informative liWM regions suggesting increased cellular infiltration and altered nuclear architecture in short survival associated areas while tiles containing few neuronal cell bodies (largest nuclei sizes) near the grey matter belonged to non-informative liWM regions.

**Figure 3:**
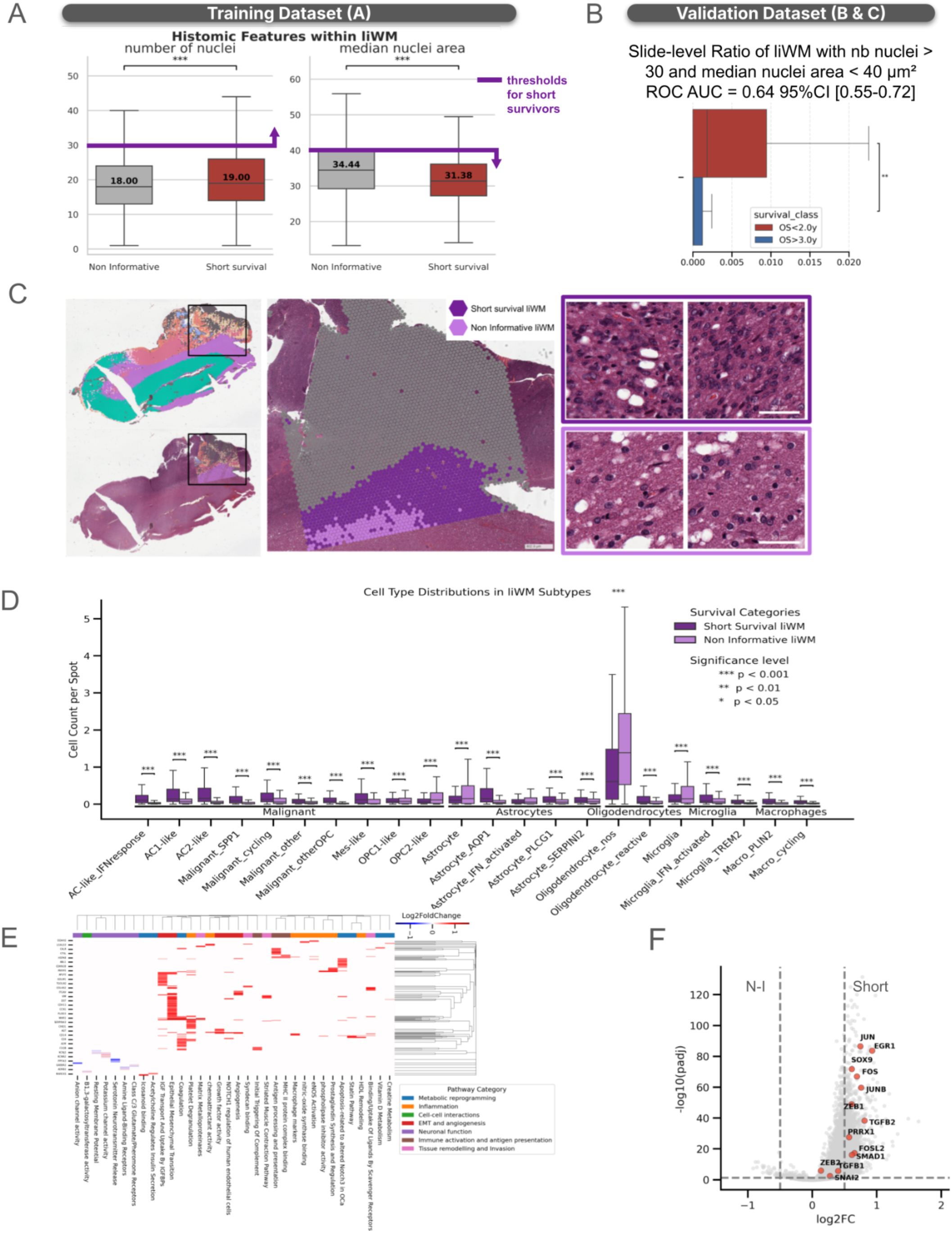
A sub-region of infiltrated white matter is predictive of short survival. **(a)** Distributions of the number of nuclei per tile and the median nuclear area per tile across 457 samples in cohort A (training dataset). Only tiles with a tile score > 0 were included. Purple lines denote thresholds optimized on the training set; arrows indicate the direction associated with shorter survival. *** indicates p value < 0.001 for a Mann Whitney U test between two boxplots. **(b)** Proportion of biomarker-positive tiles defined as infiltrated white matter (liWM) regions with >30 nuclei and a median nuclear area <40 µm² among all tumor tiles in short- and long-surviving patients from cohorts B and C (testing dataset). **(c)** Representative H&E slide and corresponding Visium spatial transcriptomics overlay (validation dataset) showing biomarker positive liWM tiles (dark purple) in comparison to adjacent liWM tissue (purple). **(d)** Cell type composition per tile in biomarker-positive liWM tiles versus non informative liWM tiles in the n=31 patients of the MOSAIC dataset. Cell types with median count per tile <0.1 were excluded for clarity. **(e)** Pathways enriched in short survival liWM vs non informative liWM. Raster plot of GSEA showing leading genes of each pathway by log2FoldChange. Only pathways with FDR<0.2 were included (See GSEA results and all genes in Supplementary table I and Table J [Supplementary material]). **(f)** Volcano plot for DEA of short survival liWM vs non informative liWM.

Based on these nuclear features, we defined a histological biomarker to identify short-survival liWM regions. Thresholds were selected to optimize predictive performance in the training dataset: a minimum of 30 nuclei per tile and a maximum median nuclear area of 40 µm². This biomarker, quantified as the percentage of tissue per slide meeting these criteria, achieved a ROC AUC of 0.64 [95% CI: 0.55–0.72] for survival prediction on the testing dataset (Fig. 3B), highlighting its generalizability across patient cohorts. The prognostic value of the slide-level mean number of nuclei per tile and the slide-level median nuclei area per tile were tested and resulted in ROC AUC values consistent with random performance.

Spatial visualization of the biomarker across H&E sections revealed consistent localization patterns, frequently concentrated at the interface between tumor core and surrounding white matter (Fig. 3C). This spatial coherence and colocalization with invasion front of the tumor supports the biological relevance of the identified histomorphological signature and its association with poor prognosis.

Second, using the SpT cohort, we compared the distribution of cell types between spots labelled short-survival versus non-informative (Fig. 3D). After excluding rare cell types (median counts per spot < 0.1), we found significant differences in cell type proportions for most cell types. Within the malignant cells compartment, and as previously mentioned and as observed in other regions, short survival spots were significantly enriched in Mes-like cells. There was also an enrichment in most other malignant cell subtypes like astrocyte-like (AC-like) subtypes and malignant cycling cells, except in oligodendrocyte progenitor cell-like type 2 cells (OPC2-like). Within the non-malignant compartment, activated astrocyte, oligodendrocyte, and microglia subtypes were enriched in short survival tiles, as opposed to non-activated subtypes in non-informative tiles. Additionally, two subsets of macrophages were also significantly enriched in short survival tiles: Macro_PLIN2 and cycling macrophages. So far, these findings indicate that liWM region is predictive of poor prognosis when it is infiltrated by a higher number of malignant cells including Mes-like and *SPP1*-expressing cells and lipid-associated macrophages.

Next, to go beyond differences in cell types, we performed differential gene expression analysis and pathway analysis between the two categories of spots associated with short survival or non-informative. GSEA analysis revealed that non-informative liWM regions are significantly enriched for neuronal function, with a strong representation of pathways involved in neurotransmitter release, the maintenance of the resting membrane potential, and the activity of glutamate and pheromone receptors (Fig 3E). This finding aligns with the observed cellular distribution in these tiles (Fig 3C). In stark contrast, short-survival liWM tiles presented a highly active and aggressive cancer-evolving microenvironment. This is characterized by an inflammatory and immunosuppressive state driven by a significant shift in lipid metabolism and extensive tumor remodeling (Fig 3E). This aggressive phenotype is further supported by the enrichment of actionable pathways related to tissue remodeling and invasion (Comba et al. 2022) (Fig 3E, and Supp Fig 10), evidenced by the co-expression of collagens (e.g., *COL1A1, COL1A2, COL5A2*) with matrix-degrading enzymes (*MMPs, TIMP2, TIMP3*), which supports active matrix remodeling required for tumor invasion in these short-survival associated liWM regions (Supp Fig 10). Moreover, there was a pronounced Mes-like/EMT signature (Fig 3E-F), sustaining the invasion front through the upregulation of key transcription factors that triggered Mes-like cell fate such as *ZEB1*, *SOX9*, and *PRRX1* (Wellner et al. 2009; Siebzehnrubl et al. 2013; Cheung et al. 2005; Sardar et al. 2022; Youssef et al. 2024; Wu et al. 2025). Our DEG analysis also uncovered an active transcriptional program promoting dedifferentiation and wound healing (Fig 3F), with high expression of genes like *JUNB*, *FOS*, and *EGR1* (Marques et al. 2021; Abou-Antoun et al. 2025; DU et al. 2023; Li et al. 2025), suggesting an active cellular plasticity promoting GBM tumor cells to transition to a more stem-like state, which may contribute to the adaptability of aggressive tumors along the invasion front.

### Long-survivor-associated tumor regions exhibit enhanced angiogenesis and endothelial Tip Cell activity

Our tile prediction model revealed an enrichment of long-survivor tiles in the Tumor 1.0, Tumor 1.1, and Tumor 2.1 regions (Fig 2E). To further analyse the features and underlying biology of those tiles, a similar methodology was applied across all three tile categories: predictive of long-survival, short-survival, and non-informative (Fig 4A).

**Figure 4:**
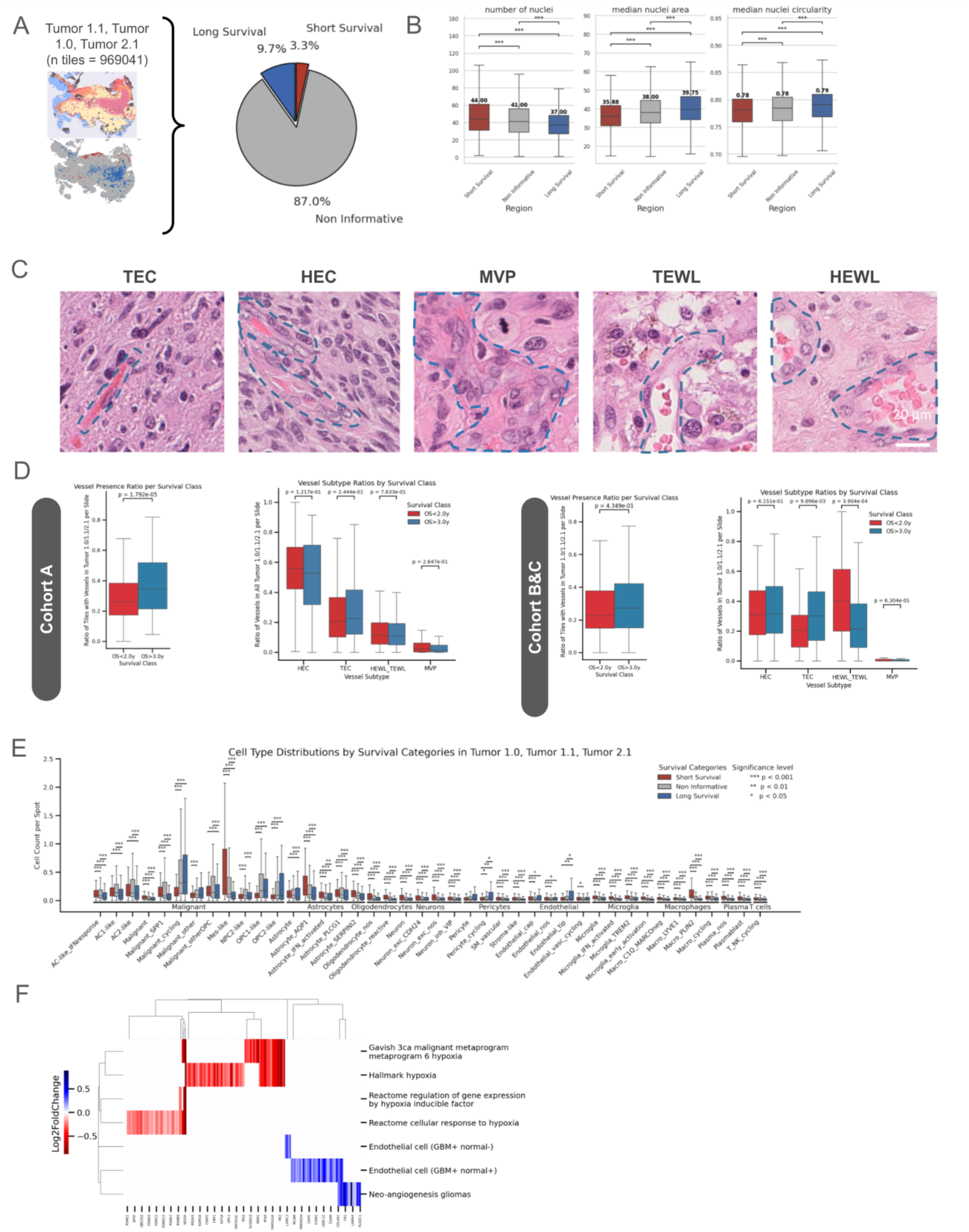
Angiogenesis is increased in areas predictive of long survival. **(a).** In cohort A, proportion of tiles associated with short survival, long survival, and non-informative categories in pooled tissue subtypes Tumor 1.0, Tumor 1.1, and Tumor 2.1. **(b).** Within pooled tissue subtypes Tumor 1.0, Tumor 1.1, and Tumor 2.1, distributions of nuclear morphology features like nuclear density (number of nuclei per tile) and nuclear size (median nuclear area in µm²) and median nuclear circularity. **(c)** Representative images of the annotated vessels subtypes patterns on H&E slides (20X): (i) thin endothelium capillary (TEC), (ii) hyperplasic endothelium capillary (HEC), (iii) Microvascular Proliferation (MVP) (iv) Thin Endothelium Wide Lumen (TEWL) (v) Hyperplasic Endothelium Wide Lumen (HEWL) **(d)** Relative ratios of tiles with vessels or specific vessel subtypes within tumor regions (1.0, 1.1 and 2.1) for each patient across the two survival groups (2y < OS and OS > 3y). Distributions are computed for the training set (cohort A) and validation set (cohort B&C). Vessel presence and vessel subtypes detection are obtained after inference by the vessels detection and vessels subtypes classifier models on the entire cohorts. **(e)** Distribution of cell count per cell type in the tumor regions (1.0, 1.1 and 2.1) for short-survival, long-survival and non-informative tiles. Distributions of cell counts statistically significantly different (Mann–Whitney test; threshold at 0.05) are indicated with a star. 31 samples from Cohort B with paired H&E, Visium spatial transcriptomics, and scRNAseq were used. Cell types with low range of mean count variations across tissue regions were removed for clarity. **(f)** Illustration of the pathway enrichment analyses of the differential expressed genes between long- and short-survival. Negative fold change is associated with worse survival outcomes and positive with better survival outcomes. (See GSEA results and all genes in supplementary table L [Supplementary material])

Using cohort A, we refined the histopathological characterization of these tile categories by quantifying nuclear morphology. For each tile within the H&E-stained slides, we computed (i) the number of nuclei per tile, (ii) the median nuclear area, and (iii) the median nuclear eccentricity. Notably, in all three regions, long-survival tiles exhibited significantly lower nuclear density and larger median nuclear area compared to both short-survival and non-informative tiles (p < 0.001; Fig. 4B, Supp. Fig. 11). Based on this reduced nuclear density, and supported by visual inspection of high-resolution histology tiles from Tumor 1.0, Tumor 1.1, and Tumor 2.1 (Supp. Fig. 12), we hypothesized that long-survival regions may be enriched in blood vessels, which prompted us to perform quantification. To assess whether the presence and subtype of vessels were associated with survival outcomes, we developed a model to detect the presence or absence of blood vessels within tiles and classify five vessel subtypes (Fig 4C, Supp Fig. 13): MVP, thin endothelium capillary (TEC) as the normal aspect of brain capillaries, hyperplastic endothelium capillary (HEC) with increased density and/or thickness of endothelial cells as the intermediate state of tumor neoangiogenesis between normal capillaries and MVP, and thin endothelium with wide lumen (TEWL) grouped with the hyperplastic endothelium with wide lumen (HEWL) which can correspond to veinules or dilated capillary (Daumas-Duport et al. 1997; Nalisnik et al. 2017). The model was trained and evaluated using blinded annotations from an expert neuropathologist on the training (A) and testing (B&C) cohorts (Supp. Tables G and H; <u>Methods</u>). The model was subsequently applied to detect and classify blood vessels across cohorts A, B, and C within Tumor 1.0, Tumor 1.1, and Tumor 2.1 (<u>Methods</u>). The presence and subtype distribution of vessels was then compared between end-of-spectrum survival groups (OS<2y and OS≥3y) (Fig 4D). As we hypothesised, samples from long survivor patients exhibited an overall higher prevalence of blood vessels compared to short survivor patients in the training cohort A *(p = 1.79 × 10*⁻⁵*, p < 0.001)*. A similar trend, although not significant, was observed in external validation cohorts B&C *(p = 4.35 × 10*⁻*^1^, ns)*. Regarding blood vessel subtypes, the ratio of TEC was higher in long survivors compared to short survivors (*p = 9.90 × 10*^⁻^*^3^, p < 0.01* in Cohorts B&C, *p = 2.44 × 10*^⁻^*^1^* not significant in Cohort A) while other subtype ratio differences were not significant (p≥ 0.05) (Fig 4D, Supp. Fig 11-12).

We further interrogated the cellular differences between long- and short- survivor tumor regions by analysing the cellular distribution across these tissue tiles. Long-survivor tumor tiles exhibited a higher proportion of NPC-like and OPC-like malignant cells *(p < 0.001)* and a significant enrichment of endothelial (Endothelial_nos *(p < 0.05)*, Endothelial_tip *(p < 0.01)*) and pericyte (Pericytes, Pericyte_cycling, *p < 0.05*) cells, consistent with active growing and orchestrated vessel formation (Gerhardt et al. 2003; Hellström et al. 2007; Li et al. 2024; Colwell et al. 2017) (Fig 4E). This finding is consistent with the increased angiogenesis indicated by our ML-driven H&E analysis (Fig 4B-D). In contrast, short survival tiles contained a significantly higher proportion of Mes-like malignant cells *(p < 0.001)* and reactive astrocytes (Astrocytes_AQP1, *p < 0.001*), both of which are associated with a highly invasive and hypoxic tumor microenvironment (Neftel et al. 2019; Kloosterman, Erbani, Boon, Farber, Handgraaf, Ando-Kuri, Sánchez-López, Fontein, Mertz, Nieuwland, Liu, Forn-Cuni, van der Wel, et al. 2024). We also observed a general enrichment of myeloid and lymphoid subsets, particularly some subsets of macrophages (PLIN2 and LYVE1, *p < 0.001*) which presence is associated with transition to Mes-like phenotype in hypoxic environments, resistant to therapy and remodeling (Governa et al. 2024; Miller et al. 2025; Deligne et Midwood 2021; Elfstrum et al. 2024) (Fig 4E).

Lastly, to further interrogate the differences between GBM long and short survivor categories within these tumor regions, we performed differential gene expression and pathway enrichment analyses (Fig. 4F, <u>Methods</u>). So far, we identified a differential trend in the distribution of blood vessels (Fig 4A-B) and hypoxic-associated cellular subsets across tile categories (Fig 4E), therefore we focused our analysis on pathways associated with those biological processes (Fig 4F). GSEA analysis revealed a significant enrichment in short survivor tiles of hypoxia-triggered pathways (Fig 4F). In contrast, long survivor associated tiles showed significant upregulation of neo-angiogenesis (Lothar C Dieterich et al. 2012), and significant enrichment of both tumor-specific vasculature (Dusart et al. 2019) and the pre-existing, normal vasculature (Dusart et al. 2019), suggesting an orchestrated balance of endothelial cells remodeling the existing vasculature while simultaneously stimulating the creation of new, tumor-driven vessels.

Overall, our results demonstrated that tumor regions predictive of long survival are defined by an overall increased trend of number of blood vessels and of the percentage of blood vessels belonging to the TEC subtype. These findings are supported by the significant enrichment of *de novo* vasculogenesis, in part sustained by endothelial tip cells in long survivors in contrast with an hypoxic cellular profile in short survivor tiles.

## Discussion

GBM tumors exhibit an infiltrative growth pattern, where neoplastic cells propagate through interconnected, heterogeneous microenvironments. These tumor niches are characterized by a dynamic interplay of varying oxygen tensions and diverse cellular composition (Neftel et al. 2019). Despite extensive genomic profiling of GBM patient samples, the distinct histological and molecular features that underpin end-of-spectrum long-term survival remain poorly understood. Here, we used an advanced machine learning pipeline on spatial histological patterns and transcriptomics across a new GBM dataset with the largest number or end-of-spectrum long-term survivor (OS ≥ 3 years) to date. Unique anatomical areas have emerged as novel spatial predictive biomarkers for short and long survival. We also unveil a functional angiogenesis underlying biology of long survivor tumors.

GBM is the most prevalent and aggressive brain tumor in adults, and it remains a major unmet medical need. While many studies have explored survival prediction in GBM using histology, most have relied heavily on the TCGA dataset despite its significant limitations (Redlich et al. 2024). In addition, most of those studies lacked external validation on independent datasets, as most models were trained and tested only on TCGA data. Only two studies (Redlich et al. 2024; Verma et al. 2024; Zadeh Shirazi et al. 2020) performed external validation (n=250 and n=9). This raises concerns about how well these models generalize.

The TCGA glioma cohort includes a heterogeneous mix of tumors, with only a subset (32%) that meets the current 2021 WHO diagnostic criteria for GBM (Hewitt et al. 2023). This makes it challenging to identify features predictive of long-term survival, as only a 8.5% of the cohort exhibit long-term survival (OS ≥ 3 years). Furthermore, the TCGA data span diagnoses from 1989 to 2012, meaning many patients were treated before the 2005 introduction of the Stupp protocol, the current standard of care. This therapeutic heterogeneity, combined with incomplete treatment details (Brennan et al. 2013), and poor inclusion criteria documentation (Redlich et al. 2024), introduces considerable uncertainty when linking histology to outcomes. Alternative datasets, such as CPTAC-GBM (« CPTAC-GBM » 2018), also have similar constraints. Specifically, in CPTAC-GBM, it is unknown whether tissue samples were fresh-frozen or FFPE and patient survival information is not available in other datasets (« CPTAC-GBM » 2018; Puchalski et al. 2018) which preempted their use in our work.

Predicting survival is inherently difficult due to the many variables at play, including patient comorbidities, tumor biology, and random events (Liu et al. 2023; Wang et al. 2023; Chen et al. 2021; Zhu et al. 2017). While previous studies have relied on complex multi-modal or limited uni-modal approaches (Redlich et al. 2024), we addressed these challenges by using a new dataset with well-defined inclusion criteria and minimal censorship. This allowed us to precisely define survival endpoints and ensure a sufficient number of long-term survivors for model training.

Importantly, our approach goes beyond mere survival prediction. By leveraging a tile-level prediction model (Lazard et al. 2022), we were able to link predicted survival classes (short vs. long) to specific regions within whole slide images. This allowed for the spatial interpretation of histological patterns associated with patients’ outcome, a key step toward defining the underlying biology. Our approach was validated with the task of detecting FGFR3-TACC3 fusion from histology, which benefits from established histological markers (Bielle et al. 2018), confirming that our model accurately highlights prognostic regions with biological relevance.

To our knowledge, this is also the first time that the diagnostic accuracy of FGFR3-TACC3 fusion has been systematically assessed using only H&E-stained slides, without access to molecular testing. While FGFR3-TACC3 fusions have been associated with improved survival in GBM (Di Stefano et al. 2020), they did not confound our survival prediction model as fusion-positive cases were rare in the training cohort. Importantly, the histological patterns our model used to predict the FGFR3-TACC3 fusion were distinct from those associated with survival outcomes, indicating that these features represent separate underlying biological processes. This demonstrated the model’s ability to accurately identify known histopathological hallmarks of FGFR3-TACC3 fusion from H&E tiles, confirming its strength in detecting biologically meaningful features that predict molecular status and enable interpretable spatial characterization.

Our unsupervised learning approach uncovered 14 distinct tissue regions that are both spatially consistent and shared across different samples. These regions are spatially organized into layers, transitioning from necrotic to tumor core regions, and then to lowly-infiltrated regions. This layered organization closely aligns with the 5-layer model recently published in a spatial transcriptomics study (Greenwald et al. 2024). Notably, our model identified a peri-necrotic tumor region located directly adjacent to necrosis and enriched with MES-like and inflammatory macrophages. This finding is particularly significant because this region is consistent, both in location and cellular composition, with the hypoxia-adjacent region described in the same study - a region that did not correspond to any standard histology patterns in the IVY GAP dataset (Greenwald et al. 2024). This underscores the ability of our model to identify biologically relevant features that are not easily captured by conventional histological annotation.

We identified a novel imaging biomarker indicative of poor prognosis, the liWM region composed of areas of white matter with low tumor infiltration. This histological aspect was previously correlated with the peritumoral non contrast enhancing zone observed on presurgical MRI (Wang et al. 2024). This region is typically not targeted by maximal safe resection of GBM according to the SOC although the debated concept of a supramarginal resection targeting this tumor component has emerged to encompass it (Chang et al. 2017; Molinaro et al. 2020; Jackson et al. 2020). It harbors early tumor subclones in comparison to the tumor bulk (Spiteri et al. 2019) and it is the location of 90% of GBM relapses (Lemée et al. 2015).

White matter-infiltrating tumor cells are farther from the tumor core and may reside in regions with lower vascularity and maintained blood-brain-barrier, reducing their exposure to systemic therapies like temozolomide. The increased presence of this infiltrating component on histology may be associated with a higher burden of residual disease after resection and with tumor spread farther from the operative margins and beyond the volume targeted by irradiation. This “sanctuary” effect could allow these cells to survive initial treatment and contribute to recurrence, worsening prognosis (Ahmed et al. 2023; Oberoi et al. 2016). Additionally, the hypoxic and nutrient-poor conditions in white matter may select for more resilient tumor subpopulations, further driving resistance. As this infiltrating zone is enriched with early subclones during tumor evolution (Spiteri et al. 2019), it may contain a reservoir of fitness to cope with the pressure of selection by the treatments and reprogrammed metabolism. Infiltration into white matter could also disrupt critical neural connectivity, leading to more severe neurological deficits at diagnosis. This functional impairment might indirectly contribute to worse survival by limiting patients’ performance status and tolerance to aggressive treatments (Duffau 2014).

White matter tracts serve as “highways” for GBM cell migration due to their aligned structure and the presence of axons (Kang et al. 2025; Rao et al. 2013). Tumor cells infiltrating these regions may reflect a more aggressive phenotype with heightened migratory and invasive capabilities. Studies have shown that GBM cells exploit white matter tracts for distant spread, which could explain poorer survival due to increased likelihood of recurrence or multifocal disease (Mickevicius et al. 2015). In contrast, other studies identified pro-oligodendrocyte differentiation signals from the white matter environment slowing down tumor cells invasion and proliferation (Brooks et al. 2021). Of note, our data demonstrated that within the white matter there are informative niches associated with short survival, significantly enriched in Astrocyte-like tumor cells in contrast with non-informative regions enriched in oligodendrocytes-like malignant cells, concealing previous contrasting reports. Our pathway analysis revealed high tissue remodeling and lipid-shifting microenvironment in the short survival tiles from liWM, in part driven by PI3K/AKT signaling pathway and extracellular matrix activation of integrin signaling to facilitate invasion. These pathways have been shown to be crucial for the interaction between tumor cells and the white matter microenvironment, including oligodendrocytes and astrocytes, and promoting survival and resistance to therapy (Giese et al. 2003).

The short-survival liWM tiles presented some additional unique cellular populations compared to non-informative counterparts, including significant enrichment of Mes-like and *SPP1*-expressing malignant cells, and non-tumoral Astrocytes-AQP1 and lipid-associated macrophages PLIN2, suggesting a significant metabolic crosstalk between tumor cells and the surrounding TME in the liWM that fuels invasion and tumor growth. Indeed, recent findings indicate that *PRRX1* functions as a pioneer factor, triggering an embryonic-like EMT program that specifically sustains a cancer invasive trajectory and metastatic progression by stabilizing the mesenchymal phenotype in migratory cells (Youssef et al. 2024). In contrast, upregulation of transcription regulators *EGR1* and *JUNB* suggests a simultaneous activation of an injury-like Mes-like state (Molaei et al. 2025; Miller et al. 2025). Therefore, in contrast with Mes-like observed in the tumor core, liWM Mes-like state is characterised by an intermediate EMT program involving a hybrid transcriptional program, similar to the one identified in therapy-resistant, and progenitor-like cells and associated with poorer prognosis (Guo et al. 2025; Liao et Yang 2020; Aggarwal et al. 2021).

Astrocytes-AQP1 has been shown to promote a reactive, inflammatory environment that in turns induce a Mes-like state in tumor cells (Governa et al. 2024), while lipid-associated PLIN2-macrophages provide direct metabolic support, especially in regions like the white matter of myelin degradation, where they scavenge lipid-rich debris to supply the necessary lipids to Mes-like cancer cells(Kloosterman, Erbani, Boon, Farber, Handgraaf, Ando-Kuri, Sánchez-López, Fontein, Mertz, Nieuwland, Liu, Forn-Cuni, Wel, et al. 2024; Miller et al. 2025). Axonal injury in the infiltrated white matter promotes GBM progression through activation of neuro-inflammation (Clements et al. 2025).Furthermore, PLIN2-macrophages from liWM showed an activated lipid-processing immunosuppressive state denoted by the TREM2-TYROBP network (Read et al. 2024; Yan et al. 2024; Segura-Collar et al. 2025; Wischnewski et al. 2025)(REFS), which is not observed in the other tumor regions. Our liWM data suggest that the contribution of each of those cell types to the MES-like fate is complementary, and promotes a novel liWM Mes-like state, characterised by an intermediate EMT program involving both upregulation of invasion (Wellner et al. 2009; Siebzehnrubl et al. 2013; Cheung et al. 2005; Sardar et al. 2022; Youssef et al. 2024) and inflammation programs (Marques et al. 2021; Abou-Antoun et al. 2025; DU et al. 2023; Li et al. 2025). This may be creating a positive feedback loop between liWM Mes-like, PLIN2-mac and astrocytes-AQP1, as well as defining functional dependencies to explore therapies.

In the tumor core regions, our ML-driven method identified some areas of tumor tissue highly predictive of better prognosis (long-survivor tiles). These long-survivor tumor tiles localise at an intermediate position between tumor core and infiltrating zone (liWM), far from the necrotic and perinecrotic regions. We observed an enhanced presence of vessels trend, a histological feature suggestive of angiogenesis (Nalisnik et al. 2017) in tumor regions associated with longer survival. This observation was consistent across all testing and validation cohorts, suggesting a potential link between increased angiogenesis and a more favorable prognosis, even if the finding did not reach statistical significance in every case. This observation was further supported by the significant enrichment of endothelial cells and pericytes in long-survivor tiles, which are crucial for vessel formation (Dusart et al. 2019; Lothar C. Dieterich et al. 2012). Specifically, the enrichment of endothelial_tip cells points to new, functional capillary growth. The enrichment of these particular cells in long-survivor tumors suggests that the type and function of the vasculature are critical, as opposed to simply its presence. The presence of a stable, functional vasculature is further supported by the downregulation of the hypoxia response signature in these tiles, which correlates with significantly lower numbers of Mes-like malignant cells and PLIN2-expressing macrophages—both of which are associated with hostile, hypoxic environments. Our data collectively suggests that efficient perfusion from this newly formed vascular network contributes to a less aggressive tumor microenvironment.

These findings challenge the conventional view that all tumor vasculature is chaotic and pro-invasive, corresponding to MVP (Nalisnik et al. 2017; Jain et al. 2007; Jain 2005; Sorensen et al. 2009). Instead, it suggests that a more organized, or ‘normalized,’ vascular network exists in less aggressive tumor areas. Therefore, while angiogenesis is a canonical hallmark of GBM (Hardee et Zagzag 2012), our results showed that its prognostic significance is more complex than previously thought. Our analysis supports that tumor regions with a better prognosis and longer patient survival exhibit a distinct type of angiogenesis. Indeed, this angiogenesis is distinct from MVP, which is often located closer to the tumor core and less efficient for perfusion. Such a functional angiogenesis producing capillaries rather than MVP in the tumor intermediate zone could improve patients outcome through several mechanisms: better diffusion of chemotherapy inside the residual tumor, increased effect of radiotherapy due to higher oxygen pressure, decrease of glioblastoma stem cell state related to hypoxic niche, and endothelium more functional for extravasation of antitumoral immune cells (Markwell et al. 2022; Jain et al. 2007; Jordan et Sonveaux 2012; Beckers et al. 2024; Colwell et al. 2017). Further investigations are required to better understand the cellular and metabolic ecosystem related to angiogenesis-related improved prognosis.

Overall, our findings challenge prior assumptions about GBM heterogeneity and survival prediction, highlighting how subtle histological cues can reveal underlying biological processes and functional differences in angiogenesis and tumor invasion. Ultimately, our study provides a framework for applying machine learning to uncover biologically relevant spatial biomarkers, offering new insights into GBM pathogenesis and paving the way for more precise therapies.

## Acknowledgements

We are grateful to patients, physicians, nurses and research assistants involved in the study.

We thank for their contributions Yannick Marie, Catherine Carpentier and the Onconeurotek databank (Paris Brain Institute) for data collection and molecular data acquisition, the Onconeurotek biobank (AP-HP) for DNA samples cession to perform retrospective tests, Marine Giry and Rémi Foucart for MGMT promoter pyrosequencing (Paris Brain Institute), the technical staff of the Neuropathology Department (Pitié-Salpêtrière, AP-HP) for histological methods, immunostainings, DNA extraction, and Besma Barka for digitalization of histological slides and management of DNA methylation profiling, HISTOMIC platform for access to pathology slide scanners, Karima Mokhtari for prospective validation and collection of pathology diagnosis (Pitié-Salpêtrière, AP-HP), Hélène Madry and Badreddine Mohand Oumoussa at the Plateforme Post-génomique de la Pitié Salpétrière P3S (Sorbonne Université) for DNA methylation profiling, Thomas Gareau (Data Analysis Core, Paris Brain Institute) for the processing of DNA targeted NGS data and DNA methylation data.

We thank the Translational Research Unit, Institute of Tissue Medicine and Pathology, University of Bern, for their valuable support in retrieving the samples from the block archives and scanning them.

We thank Pr Samir Jabari and the Neuropathology Department of Universitätsklinikum Erlangen for their contribution.

This study also makes use of data generated by the MOSAIC consortium (Owkin; Charité – Universitätsmedizin Berlin (DE); Lausanne University Hospital - CHUV (CH); Universitätsklinikum Erlangen (DE); Institut Gustave Roussy (FR); University of Pittsburgh (USA)), glioblastoma cohort of the MOSAIC consortium, a non-interventional clinical trial registered under NCT06625203.

## Methods

### Dataset

This study was based on four datasets from center A (Hôpital Universitaire de la Pitié Salpêtrière AP-HP, Paris, FR), center B (Universitätsklinikum Erlangen, DE), center C (Insel Gruppe AG, Bern, CH) and a glioblastoma cohort of the MOSAIC consortium, a non-interventional clinical trial registered under NCT06625203.

#### Inclusion criteria

Patients with initial surgery (resection or biopsy) of GBM performed between 2005 and 2018. were retrospectively included after review by a senior neuropathologist (FB). The diagnostic was updated according to WHO classification 2021 5th edition. Only samples containing necrosis and/or MVP on digitalized H&E slides were considered. Above age of 55 years, negative IDH1 R132H immunostaining and maintained expression of ATRX were required whereas under age of 55 years, IDH1 and IDH2 wildtype status was required. Infratentorial location, relapsing tumor, alternative tumor type at retrospective review, ATRX loss of expression, presence of pathogenic mutation of IDH1, IDH2 or H3F3A, loss of H3K27me3 staining (for tumor infiltrating the midline) were exclusion criteria. Cohort A was also enriched with GBM bearing the FGFR3-TACC3 gene fusion, which has been also associated with better outcome (Di Stefano et al. 2020).

#### Methylation profiling

Whole-genome methylation profiling was performed using Infinium MethylationEPICv2 following manufacturer’s instructions after DNA extraction from tumor samples. Tumor classification was predicted from the Molecular Neuropathology methylation profiling classifier v12.8 (Capper et al. 2018).

As the essential diagnostic criteria of GBM according to WHO classification may lack specificity and may overlap with other gliomas of better prognosis, we performed methylation profiling of 50 GBM with OS > 5y of the training set. We observed that 45 matched with the GBM methylation class, one matched with “Adult-type diffuse high grade glioma, IDH-wildtype, subtype E” methylation class and 4 were inconclusive confirming the high specificity of our diagnostic review to include long survivor GBM patients.

#### Patient inclusion and FGFR3-TACC3 fusion testing

FGFR3-TACC3 fusion status was available exclusively for Cohort A (Supp. Table C). A total of 440 patients were included, of whom 44 were fusion-positive. Patient inclusion and testing FGFR3-TACC3 fusion status were obtained by first testing of FGFR3 protein expression through IHC, followed for positive cases by molecular confirmation: (i) reverse transcription and PCR of the fusion breakpoint followed by sequencing of the PCR products or (ii) RNASeq panel FusionPlex® (Integrated DNA Technologies, Archer).

#### H&E quality checks and preprocessing

We collected H&E slides of 802 patients from 3 centers. Quality control of the histological slides was conducted by two experienced pathologists (F.B. and L.C.) to ensure compliance with predefined quality criteria prior to analysis. Each slide was evaluated for sufficient tumor content and absence of imaging artifacts. Specifically, a minimum of 5 mm² of tumor tissue, free from image artefacts, was required for inclusion. The artefacts considered were air bubbles, out-of-focus regions, pen marks, folded tissue, inadequate staining, tissue truncation, or scanning errors. Only slides meeting these stringent standards were retained for subsequent analyses, ensuring the reliability and integrity of the dataset. All the 748 slides used in this study passed this assessment and slides, initially considered for this study, that did not pass this assessment were not included (9 slides from Cohort A, 17 slides from Cohort B, 28 slides from Cohort C).

After quality control, the tissue segmentation of each whole slide image is obtained automatically using a UNet convolutional neural network, previously trained on a large multi-organ dataset of whole slide images to segment the tissue regions free of imaging artefacts. The tissue regions of each whole slide image is then tessellated into tiles of size 224x224 pixels at a resolution of 0.5 micron per pixels and each tile is transformed into an embedding vector of 1536 features using the histology foundation model H-optimus-0.

#### TCGA cohort pre-processing

We began with the full TCGA glioblastoma (GBM) cohort, comprising *N* patients. To ensure molecular homogeneity and relevance to IDH–wild-type GBM, we excluded all cases harboring IDH1 or IDH2 mutations, consistent with the current WHO classification. Additionally, patients treated (strictly) before 2005 were removed. In addition, patients whose histology slides failed our predefined quality control criteria were removed from further consideration. After applying these filters, a total of 159 patients remained and were included in the downstream analyses.

### Quantification of nuclear morphology with HistoPLUS AI model

To enable large-scale, standardized quantification of nuclear morphology across whole-slide images, we deployed HistoPLUS model (Adjadj et al. 2025; Bannier et al. 2025), an AI-based tool designed for automated detection, segmentation, and classification of cells on hematoxylin and eosin (H&E)-stained slides. HistoPlus is built upon CellViT (Hörst et al. 2024), a Vision Transformer–based architecture tailored for histopathological image analysis. The model was originally trained on a dataset comprising 1,479 image patches (each 112 μm × 112 μm) derived from H&E-stained TCGA slides across five cancer types: bladder cancer (n=400), colon adenocarcinoma (n=242), lung adenocarcinoma (n=231), lung squamous cell carcinoma (n=220), and mesothelioma (n=386).

Given that brain-specific cell types were not included in the training data, we used HistoPLUS solely for nuclear detection and segmentation, without leveraging its classification outputs. This ensured unbiased quantification of nuclear features such as count, area, and eccentricity across all tiles in the GBM H&E slides.

### Tissue regions characterization

#### Identification of tissue regions on H&E slides with clustering

The tissue clustering was obtained using K-means clustering of the H-optimus-0 tile embeddings on the training cohort using a subsample of 1000 tile feature vectors per slide. The optimal number of clusters (K) was determined using a combination of methods including measure of Silhouette Scores for several values of K, hierarchical clustering and measure of Adjusted Rand Index (ARI) as illustrated in Supp Fig 4. A higher ARI indicates a higher similarity between clusterings. Jumps of ARI values are observed at k=2,3,4,5,7,9,11 and 14. Which we interpret as the numbers of clusters which introduce additional biological information in the clustering. We chose the optimal number of clusters as 14, which marked the last jump of ARI values inside the interval of maximal silhouette score.

The centroids of each cluster obtained on the training cohort were saved and were used to assign tiles of the testing dataset to clusters using a nearest neighbour approach.

Sampling of most representative tiles for each cluster (Supp Fig 5) was obtained by identifying the tiles with the smallest euclidean distance to the cluster’s centroids in the feature space.

The clusters obtained on the training dataset were annotated by pathologists (F.B. and L.C.). Annotations include the nature of the tissue (tumor, necrosis, or hemorrhage) and an assessment of tumoral cells and oedema presence and density.

Identification of tumor tissue of the whole slide image is obtained by grouping together all tumor regions: liGM, liWM, Tumor 1.0-4.0, rare tumor phenotypes, and perinecrotic tumor.

#### Tissue regions characterization via pathologist validation

To validate the tissue subtype labels derived from our unsupervised clustering approach, a board-certified neuropathologist (FB), blinded to model outputs, annotated 198 H&E slides from cohort A with the estimated percentage of five histological categories at the slide level: tumor, necrosis, healthy or lowly-infiltrated tissue, hemorrhage, and artifact.

We then computed Pearson correlations between the annotated tissue proportions and the fraction of tiles assigned to the corresponding predicted classes: Tumor: correlation with tiles labeled as liWM, Tumor 1.0–4.0, or Perinecrotic Tumor; Necrosis: Necrosis and Pre-necrosis tiles; Healthy tissue: liGM tiles; Hemorrhage: Hemorrhage tiles.

We observed strong correlations for tumor (r = 0.74), necrosis (r = 0.82), healthy tissue (r = 0.73), and hemorrhage (r = 0.54), all with p < 0.001, supporting the validity of the predicted tissue subtype annotations.

#### Tissue subtype characterization via nuclear morphology and density

To further characterize the tissue regions identified by our unsupervised clustering, we quantified nuclear features across all H&E tiles using segmentation masks generated by the HistoPLUS model. For each tile, we computed: (i) the number of nuclei, (ii) the median nuclear area, (iii) the nuclear density (defined as the ratio of total nuclear surface to tile surface), and (iv) the median nuclear eccentricity (Supp. Fig. 6).

The distributions of these features were compared across all tissue regions in both the training and testing datasets (Supp. Fig. 6). The ranking of regions by nuclear features was highly consistent between datasets and aligned with expert pathological interpretation (Table 2). As expected, liGM and liWM, annotated as low-infiltrated regions, exhibited the low nuclear density, while tumor regions showed a gradient of increasing density, peaking in Tumor 2.0, Tumor 2.1, and Rare tumor phenotypes. The Necrosis region had the lowest nuclear density overall.

liGM and Tumor 1.1 exhibited larger median nuclear areas, consistent with the presence of large neurones, as noted by the pathologist. In contrast, Tumor 3.0 and Tumor 4.0 exhibited the smallest median nuclear areas among tumor subtypes.

Analysis of nuclear eccentricity further supported subtype interpretation: liGM showed the highest nuclear circularity, consistent with enrichment in oligodendrocytes, which are characteristically round-shaped (Nestor 2008). Tumor 4.0, by contrast, had the lowest circularity among tumor regions.

Differences between the training and testing datasets were primarily observed in the relative frequencies of tissue subtypes (Supp. Fig. 6), rather than in their nuclear feature distributions. Specifically, the testing dataset contained a higher proportion of tiles classified as Artefact, Tumor 1.1, and Tumor 4.0, while the training dataset had more tiles assigned to Rare tumor phenotypes, Tumor 2.1, and Hemorrhage. These differences reflect the unsupervised clustering procedure (k-means) used to define tissue subtypes in the training cohort. As clustering aimed to create subtype regions of approximately equal tile counts [23], and no subtype information was used for patient selection in the test set, variation in subtype prevalence across cohorts was expected.

#### Tissue regions characterization via spatial organization

Beyond capturing distinct morphological features, the tissue regions demonstrated notable spatial organization. Each tissue region exhibited clear spatial localization, with minimal overlap between regions, as quantitatively confirmed by Moran’s I spatial autocorrelation index (Supp Fig. 6B), with median values ranging from 0.4 to 0.8. To compute Moran’s I, the cluster maps were first binarized (assigning 1 to tiles of the cluster of interest, 0 otherwise), and spatial clustering was assessed using K-nearest neighbor (k=4) spatial weights. The index was computed per cluster and per slide, considering only clusters with at least 20 tiles, and then aggregated across the dataset.

Tissue regions such as tumor-adjacent areas and necrotic zones, including liGM, liWM, necrosis, and pre-necrosis, displayed the highest Moran’s I values, indicating strong spatial coherence. In contrast, tumoral subtypes exhibited lower, though still positive, Moran’s I values, suggesting they are more spatially intertwined. Nevertheless, the consistently positive values across all clusters indicate a general spatial consistency among tissue subtypes.

This robust identification of localized morphological patterns revealed an underlying organization of tumor subregions that had not been previously identified by regular pathological evaluation. As shown in Fig. 2C, a structured spatial hierarchy was observed in a substantial proportion of samples: a tumor core composed of Tumor 2.0 or Tumor 2.1, characterized by high nuclei density, was often surrounded by a layer of cluster Tumor 1.0 or Tumor 1.1, with a lower nuclei density according to cell level features (Fig. Supp 6A). This layer was often bordered by liWM, which in turn was typically enclosed by liGM. Notably, liGM frequently displayed a convoluted morphology, consistent with the gyral patterns of cortical grey matter (Fig. 2A, 2C).

To further confirm this observation, we constructed an affinity graph between tissue regions using the Fruchterman–Reingold force-directed layout (Fruchterman et Reingold 1991), based on an affinity matrix derived from the 5%-Hausdorff distances between tissue regions within each slide. Separate affinity matrices were computed for the training and testing cohorts and combined using a weighted average based on the number of slides in each set. Edges with affinity values below 0.5 were removed for clarity (Fig. Supp 6C). The resulting graph reflected biologically plausible spatial proximities: liGM was closely associated with liWM, which in turn neighbored Tumor 1.0, followed by Tumor 1.1. Pre-necrosis and necrosis exhibited particularly high affinity, indicating consistent spatial proximity across slides, whereas liGM and necrosis appeared as the most distant pair, emphasizing their distinct spatial localizations within the tissue.

### Single-nuclei transcriptomics preparation, QC and annotation

Single-nuclei data was generated with the 10X Chromium FLEX protocol from FFPE sections as described previously (MOSAIC Consortium et Hoffmann 2025). Briefly, 2x25um curls per block were dissociated with a gentleMACS Octo Dissociator, nuclei counted with Cellexa in Europe. Pools of 16 samples were processed in one reaction, aiming for the recovery of 10,000 nuclei in each sample. Samples are sequenced with 10,000 reads per cell.

10x Genomics Cell Ranger v7.1.0 was used to demultiplex reads and generate gene expression matrices for each sample. To pass QC, samples contained at least 500 cells and more than 50% of reads mapping to the probeset. SoupX v1.5.2 (Young et Behjati 2020) was used to estimate the ambient RNA expression from empty droplets and remove cell free RNA contamination in the data, considering the first 50 principal components. Cells exceeding 10% mitochondrial, ribosomal, or hemoglobin gene content, expressing fewer than 200 genes, or expressing more than 6000 genes were excluded during quality control. Following this cell filtration, genes expressed in fewer than three cells across the entire sample were removed to mitigate potential technical artifacts. Subsequently, scDblFinder v1.4.0 (Germain et al. 2022) with default parameters (19 principal components) was employed to identify and remove multiplet cells.

To perform cell type annotation, individual single-cell samples were integrated at the dataset level, leveraging scVI for alignment (Lopez et al. 2018) and scANVI (Xu et al. 2021) for transferring annotations from a predefined reference dataset. This reference dataset, comprising all MOSAIC GBM samples, underwent scaling and clustering using Uniform Manifold Approximation and Projection (UMAP), followed by manual annotation of major cell types via the cellxgene app (Megill et al. 2021) and established biomarkers. For each major cell type, a subsequent Seurat clustering subdivided cells into unique transcriptomic profiles, which were then annotated using known or novel markers. A similar Seurat clustering was applied to the malignant cell cluster to identify malignant cells’ subtypes, most of which exhibited strong correlations with previously described subtypes based on gene signatures (Neftel et al. 2019; Jong et al. 2025). The most granular cell types were designated as level 4, and these level 4 cells were subsequently organized hierarchically into broader parental categories (levels 1 to 3).

### Spatial Transcriptomics preparation and QC

Detailed methodology for the 31 MOSAIC samples used are in this reference (MOSAIC Consortium et Hoffmann 2025). In summary, 5000 spots with 55 µm resolution were profiled from a total capture area of 6.5 x 6.5 mm of FFPE samples. The capture area was selected by pathologists according to the following criteria: minimal tumor burden of 40% and covering at least 20% of the selected area as priority, followed by adjacent non-tumoral tissue, invasive margin and/or TLS if possible. Samples were sequenced at 25,000 reads per spot and repeated when saturation fell below 50%.

### Cell type deconvolution

The signature matrix at level 4, which was used for downstream deconvolution, was generated using the Seurat FindMarkers function (with default parameters) and the kappa function. This involved differential expression analysis (DEA) between sub-populations of cells and minimization of the condition number. The procedure was performed in a two step approach: i) a first DEA was conducted between cell types at level 2 (low granularity) to capture canonical markers of cell types (e.g. immune cells vs malignant cells) followed by ii) a second DEA between cell types within each group at level 2 (e.g. AC1-like vs AC2-like in the malignant cell group). All genes differentially expressed (based on adjusted p-value < 0.05) are then concatenated and a filtering procedure allowed to keep the signature matrix at optimal size (n=70 marker genes at level 2, n=100 marker genes at level 4, for each cell type). The values in the signature matrix represented the average expression of those genes in each cell type.

### Tissue regions characterization via Omics profiling

To further investigate our findings, we used data of 31 GBM patients from the MOSAIC consortium that have paired histology slide, Spatial transcriptomics (SpT, by 10X Visium SD), single-nuclei transcriptomics (SnRNAseq) and clinical data from MOSAIC (MOSAIC Consortium et Hoffmann 2025). Using spatially aligned histology slides and SpT, we considered histology tiles centred at each spot. We first predicted the tissue cluster associated with each of those spots.The snRNAseq data were annotated for 41 cell types. Using a deconvolution algorithm Cell2location (Kleshchevnikov et al. 2022) we estimated the proportion of cells of each cell type in each spot. This allowed us to study the average relative abundance of the cell types in the different tissue regions (See Fig. Supp 14).

### Machine learning models for Overall Survival prediction

#### Weakly Supervised Slide-Level Classification with H&E MIL Model

We used an attention-based MIL architecture (Ilse et al. 2018) to perform slide level classification. The model operates on pre-extracted tile-level features. Each slide is represented as a bag of tiles, with the architecture designed to aggregate tile information into a single slide-level prediction.

Initially, each tile feature vector was passed through a single-layer MLP with 256 output dimensions, applied independently across all tiles. In parallel, a separate MLP (output dimension 256), followed by a linear transformation to a scalar (dimension 1), was applied to the tile features to compute an unnormalized attention score for each tile. These scores were transformed via a softmax operation to produce tile-level attention weights. The tile features were then aggregated using a weighted sum, where the weights corresponded to the attention scores. This yielded a single slide-level feature vector of dimension 256.

This vector was passed through a second MLP with two layers (output dimensions: 256 and 128), followed by a final linear layer producing a scalar output. A sigmoid activation function was applied to the output during inference to yield a probability. Sigmoid activations were also used in all MLP layers. The implementation was developed in PyTorch v2.3.1.

To obtain interpretable tile-level scores, we defined the attention score as the output of the scoring MLP prior to the softmax operation, avoiding the dependency of tile-level values on the number of tiles per slide.

Training was conducted using a weighted cross-entropy loss, with class weights inversely proportional to class frequencies to address class imbalance. Tiles labeled as necrosis, pre-necrosis, hemorrhage, or artifact were excluded during training. Due to memory constraints associated with large WSI processing, we randomly sampled 2,000 tiles per slide during training. At inference time, all remaining non-filtered tiles were used.

We used the Adam optimizer (batch size: 64; learning rate: 1e-4; β₁=0.95; β₂=0.999; ε=1e−8; no weight decay). To improve model robustness, we trained an ensemble of 15 MIL models, each initialized with different random seeds and validation splits. Final predictions were obtained by averaging the outputs of the 15 models.

### Interpretable Tile-Level Prediction with the H&E Tile Predictions Model

To enable interpretable, tile-level analysis of survival-associated regions in H&E slides, we derived a variant of the H&E MIL model by inverting the order of the prediction and attention aggregation layers at inference. This architecture, referred to as the H&E tile predictions model, follows a method previously proposed in (Lazard et al. 2022), and provides two distinct outputs for each tile: a tile score, reflecting its contribution to the patient-level prediction, and a tile prediction, indicating the tile’s predicted class association (e.g. long vs. short survival).

We applied the inference-time modification to the ensemble of 15 H&E MIL models described previously. For each tile, we computed both the tile score and tile prediction from all 15 models, and then averaged them to obtain final, ensemble-level outputs.

We used tile scores (indicating importance for patient-level survival prediction) and a tile predictions (indicating association with long or short survival) to segment each H&E image into three distinct regions: (1) a short survival region (high tile scores, low tile prediction), (2) a long survival prediction region (high tile scores, high tile prediction), and (3) a non-informative tissue region (low tile scores). This required setting thresholds for the tile scores and the tile predictions.

This segmentation required setting thresholds for both tile scores and tile predictions. The threshold for tile scores was selected by evaluating the model’s performance on the training dataset after progressively excluding low-scoring tiles (Supp. Fig. 14). The optimal threshold, defined as the point beyond which performance declined, was –2, and this value generalized well across external validation cohorts B and C for the survival classification task as well as for the FGFR-TACC3 fusion prediction task. Notably, at this threshold, the selected tiles accounted for approximately 60% of the total attention weight assigned by the model across each slide, on average (Supp. Fig. 14).

For tile predictions, thresholds were set using empirical cutoffs at the 10th and 90th percentiles of tile prediction distributions within the training cohort. These were fixed across all datasets, resulting in thresholds of –1.7 (low prediction) and 0.0 (high prediction). This consistent approach allowed robust and interpretable spatial mapping of prognostic histological patterns across cohorts.

#### Clinical model

As a baseline for comparison with histology-based models, we developed a clinical model to predict whether patients survived less than 2 years or more than 3 years. The model consisted of an ensemble of logistic regression classifiers, trained on the same Cohort A patient cohort used for the H&E models (Fig. 1). Input features included age, MGMT promoter methylation status (assessed by pyrosequencing), biological sex, Karnofsky Performance Status (KPS), and extent of surgery (encoded as: 0 = no surgery, 1 = partial resection, 2 = total resection).

To ensure robustness, we trained 15 logistic regression models using 5-fold cross-validation with 3 repeats, and aggregated their predictions by averaging the output probabilities. Training was performed using the scikit-learn implementation (v1.6.1) with L2 regularization (penalty weight = 1) and the L-BFGS solver. To address class imbalance between the two survival categories (OS<2y vs. OS≥3y), we applied inverse class frequency weighting during loss computation.

Missing data were handled via imputation: continuous variables (e.g., age, KPS) were imputed using the median, while categorical variables (e.g., MGMT status, surgery) were imputed using the most frequent value, with all imputation based on the training dataset. Notably, KPS was missing in the testing dataset, and was imputed using the median KPS value from the training cohort.

We also evaluated an XGBoost classifier, but it showed no performance gain over logistic regression in cross-validation on the training set. Consequently, logistic regression was selected as the final baseline model for clinical prediction.

#### Survival prediction evaluation

To evaluate model performance in the survival classification task (OS<2y vs. OS≥3y), we computed both ROC AUC and PR AUC, using the model predicted probabilities and the corresponding binary survival labels used during training (class 0: OS<2 y, class 1: OS≥3y).

The C-index values were computed using the model predicted probabilities and the OS values while properly accounting for censoring status. Importantly, for the C-index computation, we included all patients, including those with OS between 2 and 3 years, providing a continuous survival assessment beyond the binary classification setting.

### FGFR3-TACC3 fusion prediction from H&E slides to validate model’s interpretability capacity

#### Manual annotation of FGFR3-TACC3-associated histology by expert pathologist

To assess whether our tile-level prediction model captures biologically relevant morphologies, we performed blinded pathological review of model-identified regions from both fusion-positive and fusion-negative samples as FGFR3-TACC3 fusions are known to be associated with distinct histopathological features (Bielle et al. 2018) (Supplementary Fig. 3A).

A total of 82 representative regions (each consisting of 3×3 image tiles) were selected from the 440 patients. Regions were chosen based on the model’s average tile prediction scores across the regions, sampling those with extreme high tile predictions (fusion-positive) or extreme low tile predictions (fusion-negative). Regions were then randomized for blinded review. Tiles were sampled in 3×3 grids to preserve contextual architectural features and ensure representative coverage (Supplementary Fig. 3B-C).

An expert neuropathologist (FB), blinded to molecular status, assessed each region for hallmark FGFR3-TACC3-associated morphologies (Supplementary Fig. 3C).

The resulting set of annotations is described in Supplementary Table G.

#### FGFR3-TACC3 fusion predictive model and evaluation

The same methodology and same model as the one described in H&E MIL model was applied to classify H&E slides according to FGFR3-TACC3 fusion status.

Specifically, each tile from the slides was encoded into one feature vector through the H-Optimus-0 feature extractor. Clustering was applied to each feature tile, and tiles belonging to the clusters necrosis, pre-necrosis, artefact and haemorrhage were excluded. The MIL model was trained similarly to the model described in section H&E MIL model during 60 epochs. Weights proportional to FGFR3-TACC3 prevalence were applied to the cross entropy loss to mitigate class imbalance.

### Differential expression analysis

Differential expression analysis (DEA) was performed on 10x Visium spatial transcriptomics data using a 2-step approach.

First, one DEA was applied to each Visium sample using PyDESeq2 v0.5.2 (Muzellec et al. 2022) on the spots of the target and control group (see below for the groups definition per figure). At this stage, samples with less than 50 spots in either group are excluded. Second, a meta analysis is performed to aggregate sample-wise DEA output.

DEA was performed to identify genes differentially expressed between specific white matter regions (Fig. 3). Three comparisons were conducted:

1. Short survival liWM vs. Non-informative liWM to uncover gene expression differences between white matter regions associated with poor patient survival and those deemed non-informative.
2. Short survival liWM vs. Other to identify genes uniquely expressed in short survival liWM when contrasted with the broader tumor microenvironment.
3. Non informative liWM vs Other.

DEA was performed to study the transcriptomic differences between spots of different survival categories: long survival (positive class) and short survival spots (negative class) (Fig. 4). This was done specifically for spots in the regions Tumor 1.0, Tumor 1.1, and Tumor 2.1. Other spots were excluded. A meta analysis was used to aggregate the results across slides.

After DEA, gene set enrichment analysis (GSEA) was performed using the implementation of gseapy v1.1.9 (Fang et al. 2023). For the analysis in Fig 3, the pathway libraries “Reactome_2022” (Gillespie et al. 2022), “MSigDB_Hallmark_2020” (Liberzon et al. 2015), “GO_Molecular_Function_2021” (The Gene Ontology Consortium et al. 2023), “KEGG_2021_Human” (Kanehisa et al. 2020), and “WikiPathways_2019_Human” (Martens et al. 2021) were used. For the analysis in Fig. 4, a custom set of pathways was considered to study specifically blood vessels and hypoxia.

To reduce the number of pathways to interpret we implemented a three filtering methods:

1. Filtering based on FDR values in the GSEA output: we removed pathways with FDR > 0.2
2. Filtering based on number of leading genes: we remove pathways with number of leading genes lower than 3
3. Merge pathways based on leading genes set intersection: if two pathways had intersecting leading genes higher than a threshold (Jaccard index coefficient > 0.5) then we kept only the pathway with the highest absolute NES value.

### Blood vessel detection and classification

#### Annotation of blood vessels subtypes by expert pathologist

Visual inspection of regions from Cohort A that were associated with long survival predictions revealed an apparent increase in blood vessel density. To systematically investigate this observation and train a model able to detect and classify blood vessels at scale within the entire cohorts, annotation of blood vessel presence and subtype classification were performed by an expert pathologist on selected tiles. Tile selection was guided by H&E tile prediction model outputs, focusing on regions from patients with extreme predicted probabilities of long or short survival.

As with FGFR3-TACC3 fusion histology annotations, regions were sampled in standardized formats: 3x3 tile grids for Cohort A and 7-tile regions spatially centered on transcriptomics-aligned tiles for Cohort B. Regions were randomly sampled from the patients with the most extreme model predictions, independent of actual survival outcomes.

A total of X regions were selected, anonymized, randomized, and reviewed by an expert neuropathologist in a blinded fashion. For each tile within these regions, the pathologist assessed the presence or absence of predefined blood vessels subtypes. These blood vessel subtypes were defined based on prior observations from the training cohort (Cohort A) included the following: (i) thin endothelium capillary (TEC), (ii) hyperplasic endothelium capillary (HEC), (iii) microvascular proliferation (MVP) (iv) Thin Endothelium Wide Lumen and (HEWL) (v) Hyperplasic Endothelium Wide Lumen (TEWL) (Supplementary Table M).

#### Machine Learning models for blood vessel detection and subtype classification

We developed two supervised machine learning models to detect and classify blood vessels from histology tiles.The first model was trained to predict the probability that a given tile contains a blood vessel. The second model, applied only to tiles predicted to contain vessels, was trained to classify vessel-containing tiles into one of four predefined subtypes: TEC, HEC, MVP and HEWL+TEWL.

The HEWL and TEWL subtypes were merged due to the rarity of TEWL in the annotated dataset, which precluded training a separate model for this class. Supp Table M.

Both models used a logistic regression architecture, trained using annotated tiles from Cohort A (Suppl Table N) in a 5-fold cross-validation scheme with 3 repeats. Training optimization was performed using a weighted cross entropy loss with class weights inversely proportional to class frequency. L2 regularization (regularization weight C= 0.01) and the LBFGS optimizer were employed. Model development was implemented using the scikit-learn library (v1.6.1).

Alternative classifiers, including MLP and k-NN, were evaluated. However, logistic regression achieved comparable or superior performance for both tasks in cross-validation on Cohort A and was therefore selected for final implementation.

At inference, model predictions were ensembled by averaging the predicted probabilities across all 15 models trained during cross-validation (5 folds × 3 repeats), to improve robustness and generalization.

## SUPPLEMENTARY DATA

**Supplementary Table A.** Dataset description per modality.

| Cohort | Patients within inclusion criteria (n) | Patients with as least one Slide passing QC (n) | Samples: Bulk RNAseq after QC (n) | Samples: Single cell RNAseq after QC (n) | Samples: Visium spatial transcriptomics after QC (n) |
| --- | --- | --- | --- | --- | --- |
| A | 557 | 548 | - | - | - |
| B | 146 | 129 | 31 | 31 | 31 |
| C | 99 | 71 | - | - | - |
| TCGA GBM | 186 | 159 | - | - | - |

**Supplementary Table B.**
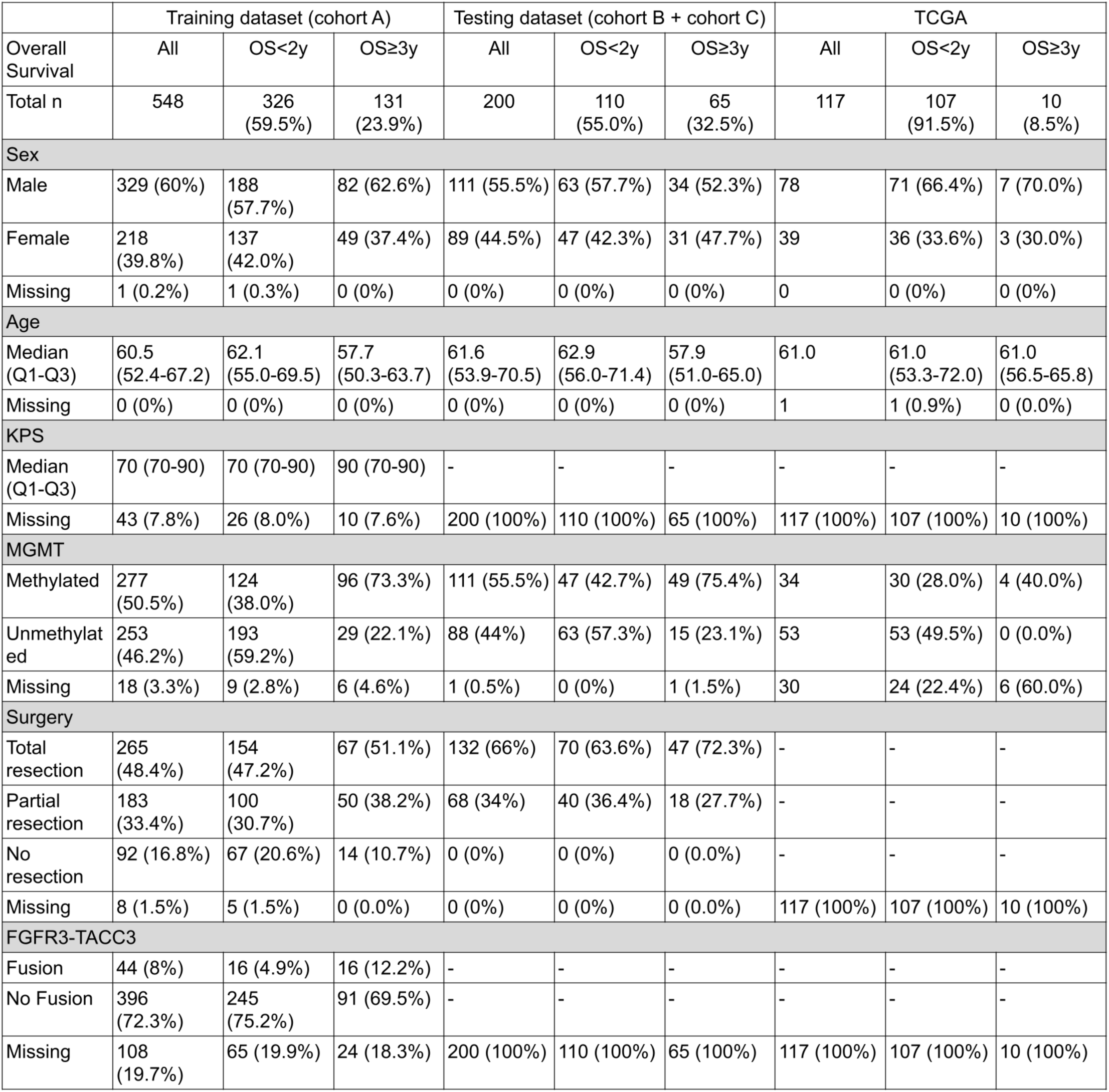
Distribution of clinical and molecular variables for the datasets used for the Survival prediction. All values are given in number with percentages in brackets, unless otherwise specified. KPS: Karnofsky Performance Status, MGMT: O6-methylguanine-DNA methyltransferase promoter.

|  | Training dataset (cohort A) |  |  | Testing dataset (cohort B + cohort C) |  |  | TCGA |  |  |
| --- | --- | --- | --- | --- | --- | --- | --- | --- | --- |
| Overall Survival | All | OS<2y | OS≥3y | All | OS<2y | OS≥3y | All | OS<2y | OS≥3y |
| Total n | 548 | 326<br>(59.5%) | 131<br>(23.9%) | 200 | 110<br>(55.0%) | 65<br>(32.5%) | 117 | 107<br>(91.5%) | 10<br>(8.5%) |
| Sex |  |  |  |  |  |  |  |  |  |
| Male | 329 (60%) | 188<br>(57.7%) | 82 (62.6%) | 111 (55.5%) | 63 (57.7%) | 34 (52.3%) | 78 | 71 (66.4%) | 7 (70.0%) |
| Female | 218<br>(39.8%) | 137<br>(42.0%) | 49 (37.4%) | 89 (44.5%) | 47 (42.3%) | 31 (47.7%) | 39 | 36 (33.6%) | 3 (30.0%) |
| Missing | 1 (0.2%) | 1 (0.3%) | 0 (0%) | 0 (0%) | 0 (0%) | 0 (0%) | 0 | 0 (0%) | 0 (0%) |
| Age |  |  |  |  |  |  |  |  |  |
| Median (Q1-Q3) | 60.5<br>(52.4-67.2) | 62.1<br>(55.0-69.5) | 57.7<br>(50.3-63.7) | 61.6<br>(53.9-70.5) | 62.9<br>(56.0-71.4) | 57.9<br>(51.0-65.0) | 61.0 | 61.0<br>(53.3-72.0) | 61.0<br>(56.5-65.8) |
| Missing | 0 (0%) | 0 (0%) | 0 (0%) | 0 (0%) | 0 (0%) | 0 (0%) | 1 | 1 (0.9%) | 0 (0.0%) |
| KPS |  |  |  |  |  |  |  |  |  |
| Median (Q1-Q3) | 70 (70-90) | 70 (70-90) | 90 (70-90) | - | - | - | - | - | - |
| Missing | 43 (7.8%) | 26 (8.0%) | 10 (7.6%) | 200 (100%) | 110 (100%) | 65 (100%) | 117 (100%) | 107 (100%) | 10 (100%) |
| MGMT |  |  |  |  |  |  |  |  |  |
| Methylated | 277<br>(50.5%) | 124<br>(38.0%) | 96 (73.3%) | 111 (55.5%) | 47 (42.7%) | 49 (75.4%) | 34 | 30 (28.0%) | 4 (40.0%) |
| Unmethylated | 253<br>(46.2%) | 193<br>(59.2%) | 29 (22.1%) | 88 (44%) | 63 (57.3%) | 15 (23.1%) | 53 | 53 (49.5%) | 0 (0.0%) |
| Missing | 18 (3.3%) | 9 (2.8%) | 6 (4.6%) | 1 (0.5%) | 0 (0%) | 1 (1.5%) | 30 | 24 (22.4%) | 6 (60.0%) |
| Surgery |  |  |  |  |  |  |  |  |  |
| Total resection | 265<br>(48.4%) | 154<br>(47.2%) | 67 (51.1%) | 132 (66%) | 70 (63.6%) | 47 (72.3%) | - | - | - |
| Partial resection | 183<br>(33.4%) | 100<br>(30.7%) | 50 (38.2%) | 68 (34%) | 40 (36.4%) | 18 (27.7%) | - | - | - |
| No resection | 92 (16.8%) | 67 (20.6%) | 14 (10.7%) | 0 (0%) | 0 (0%) | 0 (0.0%) | - | - | - |
| Missing | 8 (1.5%) | 5 (1.5%) | 0 (0.0%) | 0 (0%) | 0 (0%) | 0 (0.0%) | 117 (100%) | 107 (100%) | 10 (100%) |
| FGFR3-TACC3 |  |  |  |  |  |  |  |  |  |
| Fusion | 44 (8%) | 16 (4.9%) | 16 (12.2%) | - | - | - | - | - | - |
| No Fusion | 396<br>(72.3%) | 245<br>(75.2%) | 91 (69.5%) | - | - | - | - | - | - |
| Missing | 108<br>(19.7%) | 65 (19.9%) | 24 (18.3%) | 200 (100%) | 110 (100%) | 65 (100%) | 117 (100%) | 107 (100%) | 10 (100%) |

**Supplementary Figure 1.**
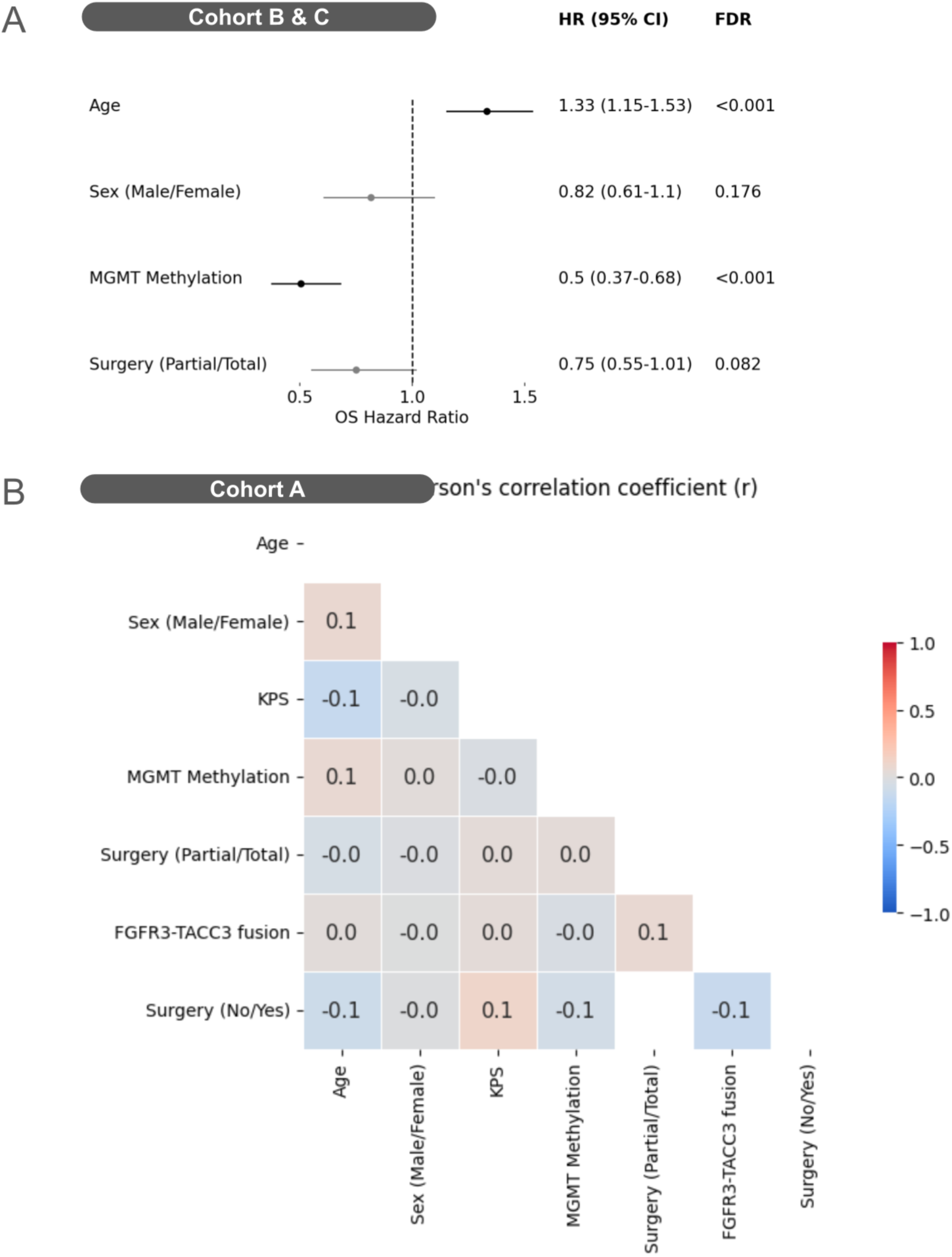
Hazard ratios of clinical variables in Cohort B and C (external validation datasets) and Pearson’s correlation between clinical variables in Cohort A (training dataset). KPS: Karnofsky Performance Status, MGMT: O6-methylguanine-DNA methyltransferase promoter.

**Supplementary Table C.** Dataset description per prediction task.

| Cohort | A | B | C | TCGA GBM |
| --- | --- | --- | --- | --- |
| Total n of pts considered | 548 | 129 | 71 | 159 |
| **End-of-spectrum survival prediction** |  |  |  |  |
| Total n of pts included | 457 (training) | 114 (external) | 61 (external) | 117 |
| N of pts with OS < 2y after QC | 326 | 83 | 27 | 107 |
| N of pts with OS ≥ 3y after QC<br>(among which n of pts with OS > 5y) | 131 (68) | 31 (14) | 34 (16) | 10 (2) |
| Total n of pts excluded | 91 | 15 | 10 | 42 |
| Patients with 2y < OS < 3y after QC (n) | 86 | 15 | 10 | 11 |
| Patients OS>2y but follow-up < 3y | 5 | 0 | 0 | 31 |
| **FGFR3–TACC3 fusion status prediction** |  |  |  |  |
| Total n of pts included | 440 | 0 | 0 | 0 |
| N of pts with fusion | 44 | - | - | - |
| N of pts without fusion | 396 | - | - | - |
| N of pts excluded for missing status | 113 | 129 | 71 |  |

**Supplementary Table D.**
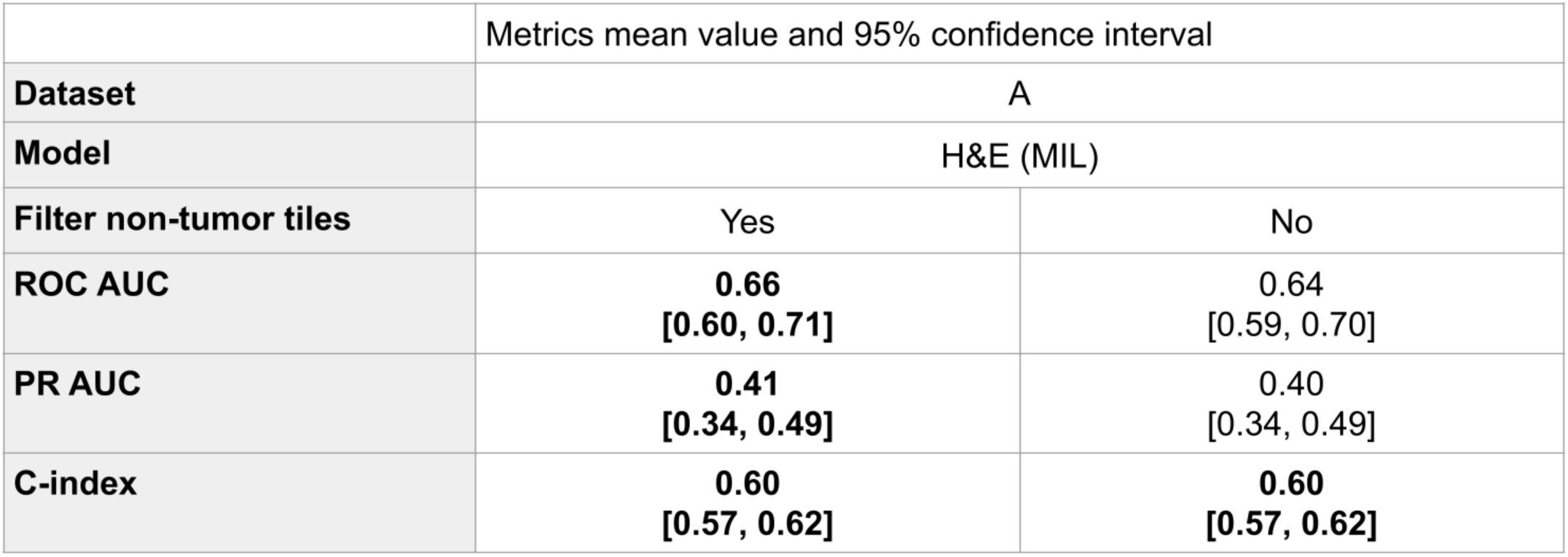
Performance of the H&E MIL model in cross validation (Cohort A) with or without filtering-out non-tumor tiles. Performance of our H&E models (ensemble of ABMIL models) with and without filtering-out non-tumor tiles using identified tissue regions. For the model with filtering, tissue regions Artefact, Hemorrhage, Necrosis, and Pre-necrosis are removed during both training and evaluation. C-index values are computed on all patients, including the ones with OS between 2 years and 3 years. Bold indicates the best mean value for each metric.

**Supplementary Table E.** Performance of the model on the training and testing cohorts on end-of-spectrum survival classification task. Performance of our H&E models (ensemble of ABMIL models), clinical model (ensemble of logistic regression using MGMT, age, surgery status, Karnofsky performance status, and gender), and the combination of H&E and clinical models on the training cohorts in cross validation and on the testing cohort to predict if a patient will either survive more than three years or less than two years after baseline diagnosis. The H&E tile predictions model is obtained by taking as prediction for each H&E slide the average of the tile predictions weighted by the softmax of the tile scores. C-index values are computed on all patients, including the one with OS between 2 years and 3 years.

|  | Metrics mean value and 95% confidence interval |  |  |  |  |  |  |  |
| --- | --- | --- | --- | --- | --- | --- | --- | --- |
| Dataset | Training cohort (Cohort A) |  |  |  | External validation cohort (Cohort B&C) |  |  |  |
| Model | Clinical | H&E (MIL) | H&E (tile predictions) | Clinical + H&E (tile predictions) | Clinical | H&E (MIL) | H&E (tile predictions) | Clinical + H&E (tile predictions) |
| ROC AUC | **0.75**<br>[0.69, 0.79] | 0.66<br>[0.60, 0.71] | 0.66<br>[0.61, 0.72] | **0.75**<br>[0.70, 0.80] | 0.76<br>[0.68, 0.83] | 0.73<br>[0.65, 0.80] | 0.73<br>[0.65, 0.81] | **0.85**<br>[0.79, 0.90] |
| PR AUC | **0.56**<br>[0.46, 0.64] | 0.41<br>[0.34, 0.49] | 0.42<br>[0.35, 0.50] | 0.53<br>[0.44, 0.63] | 0.65<br>[0.53, 0.77] | 0.66<br>[0.54, 0.76] | 0.65<br>[0.53, 0.76] | **0.77**<br>[0.65, 0.86] |
| C-index | **0.65**<br>[0.62, 0.67] | 0.60<br>[0.57, 0.62] | 0.59<br>[0.57, 0.62] | **0.65**<br>[0.62, 0.67] | 0.66<br>[0.62, 0.70] | 0.62<br>[0.58, 0.67] | 0.62<br>[0.57, 0.66] | **0.69**<br>[0.65, 0.73] |

**Supplementary Figure 2.**
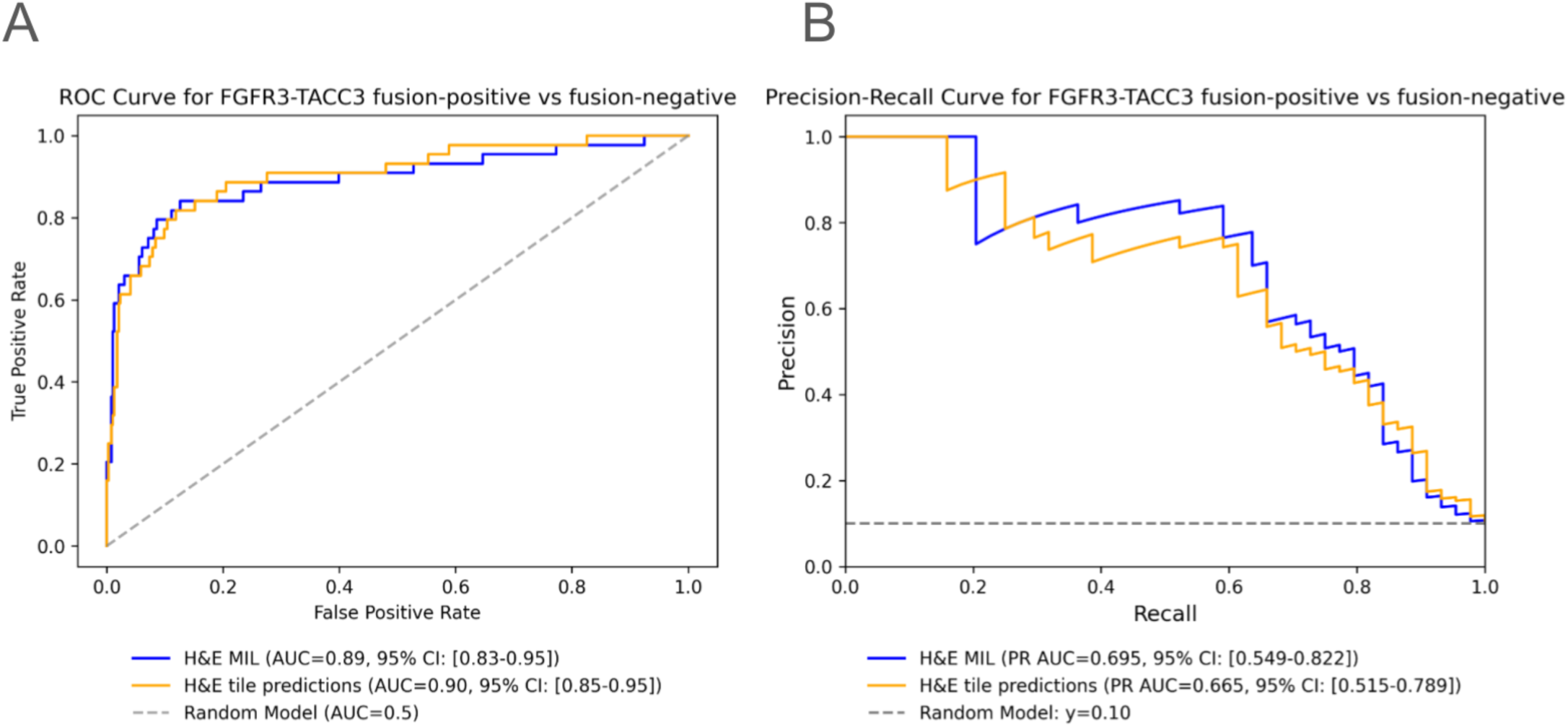
Comparison of the H&E (MIL) and H&E (tile predictions) models for the prediction of FGFR-TACC3 fusion-positive versus fusion-negative status. **(a-b)** ROC and Precision-recall curves of the H&E (MIL) and H&E (tile predictions) models to predict FGFR3-TACC3 fusion-positive or fusion-negative. The two H&E models perform similarly. All confidence intervals were obtained using bootstrapping with 1000 repeats.

**Supplementary Table F.** Performance of the model on TCGA for end-of-spectrum survival classification task. Performance of our H&E models (ensemble of ABMIL models), clinical model (ensemble of logistic regression using MGMT, age, surgery status, Karnofsky performance status, and gender), and the combination of H&E and clinical models on the training cohorts in cross validation and on the testing cohort to predict if a patient will either survive more than three years or less than two years after baseline diagnosis. The H&E (tile predictions) model is obtained by taking as prediction for each H&E slide the average of the tile predictions weighted by the softmax of the tile scores. C-index values are computed on all patients, including the ones with OS between 2 years and 3 years.

|  |  |  |  |  |
| --- | --- | --- | --- | --- |
|  | Metrics mean value and 95% confidence interval |  |  |  |
| Dataset | TCGA |  |  |  |
| Model | Clinical | H&E (MIL) | H&E (tile predictions) | Clinical + H&E (tile predictions) |
| ROC AUC | 0.55<br>[0.32, 0.75] | 0.59<br>[0.41, 0.79] | 0.60<br>[0.38, 0.82] | **0.69**<br>[0.46, 0.84] |
| PR AUC | 0.11<br>[0.04, 0.28] | 0.20<br>[0.04, 0.44] | **0.21**<br>[0.05, 0.43] | 0.14<br>[0.05, 0.28] |
| C-index | 0.58<br>[0.52, 0.64] | 0.54<br>[0.49, 0.59] | 0.54<br>[0.48, 0.60] | **0.60**<br>[0.54, 0.65] |

**Supplementary Figure 3.**
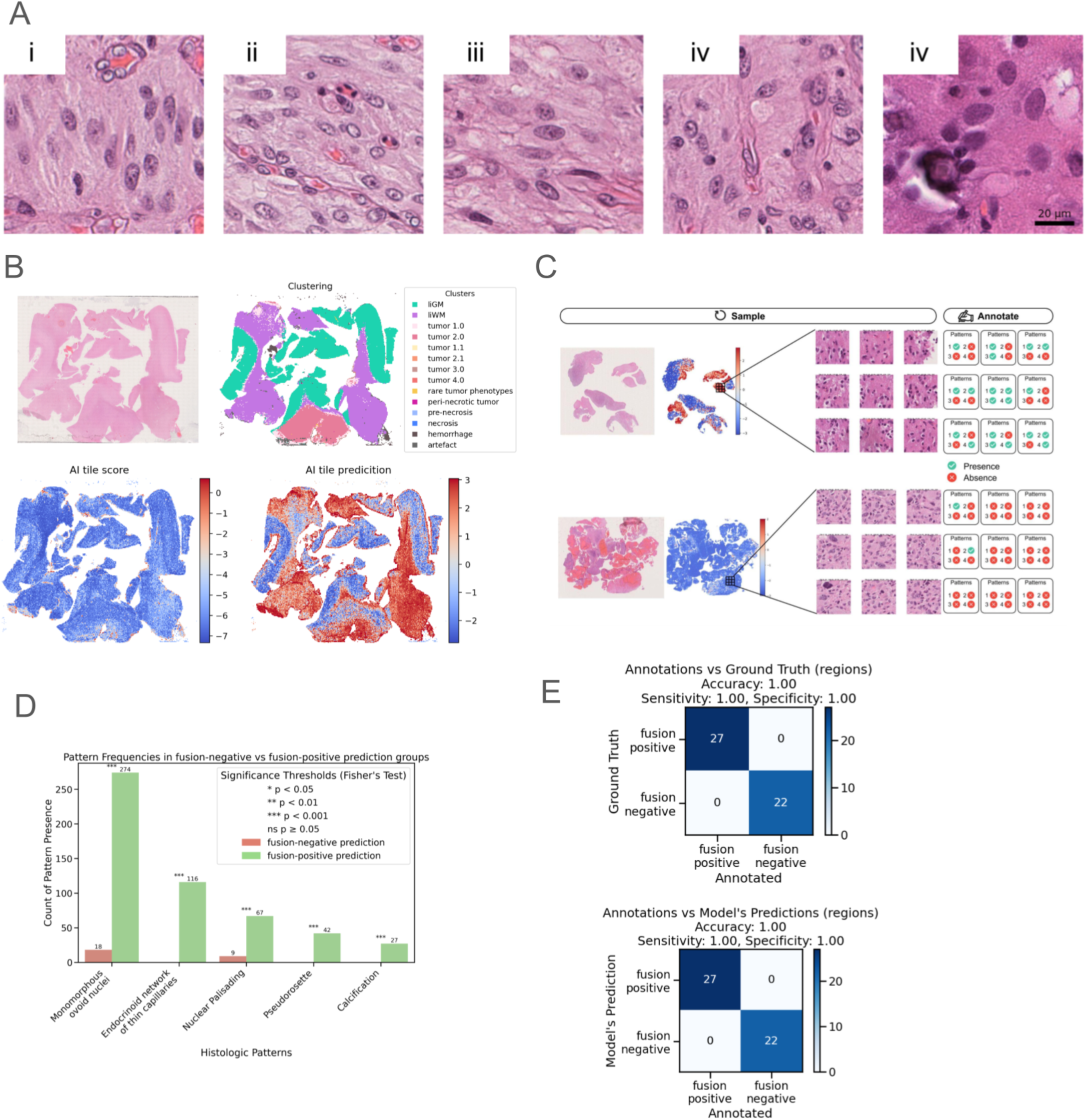
Interpretation of our model classifying FGFR3-TACC3 fusion-positive versus fusion-negative. **(a)** Illustration of five known histopathological features associated with FGFR3-TACC3 fusion and annotated by expert neuropathologist (FB). **(i)** monomorphous ovoid nuclei **(ii)** endocrinoïd network of thin capillaries **(iii)** nuclear palisading **(iv)** presence of thin cytoplasmic cell processes between tumor nuclei and vessels / pseudorosette **(v)** calcification. **(b)** Illustration of **(top-left)** FGFR3-TACC3 fusion-positive slide thumbnail **(top-right)** segmentation of tissue regions according to the clustering **(bottom-left)** heatmap of tile scores after training H&E MIL model to classify fusion-positive vs fusion-negative **(bottom-right)** heatmap of associated tile prediction. **(c)** Illustration of tile selection and annotation method: 83 representative regions (each of 9x9 tiles) were selected from both fusion-positive and fusion-negative slides according to their tile prediction scores. Regions were randomized for blinded histopathological review. Pathologist (FB), blinded for molecular status, assessed each tile for presence or absence of the five histopathological features listed in **(a)**. **(d)** Frequency of each histopathological feature across fusion-positive and fusion-negative annotated tiles. FGFR3-TACC3 fusion histological hallmarks were significantly found predominantly in tiles predicted as fusion-positive by the H&E Tile prediction model. **(e) (left)** Confusion matrix between molecular status of FGFR3-TACC3 fusion (ground truth) and pathologist estimation of the FGFR3-TACC3 fusion status performed during his blind histological review of selected regions. Confusion matrix shows 100% accuracy between pathologist annotations of known histological hallmarks of FGFR3-TACC3 fusion and the molecular ground truth, confirming the association of the patterns with the molecular status. **(right)** Confusion matrix between predictions of FGFR3-TACC3 fusion status by the of H&E Tile prediction model and pathologist estimation of the FGFR3-TACC3 fusion status performed during his blind histological review. Confusion matrix shows 100% accuracy between pathologist annotations and Tile prediction model, showing the ability of our model to focus on the same histological biomarkers as pathologists to perform its classification.

**Supplementary Table G.** Description of the Manual annotation of FGFR3-TACC3 fusion-associated histological features. This table reports the number of tiles manually annotated by an expert pathologist (FB) for morphological hallmarks associated with FGFR3TACC3 fusions. Annotated features include: monomorphous ovoid nuclei, an endocrinoid network of delicate capillaries, nuclear palisading, thin cytoplasmic processes extending between tumor nuclei and vessels or forming pseudo-rosettes, calcifications, and desmoplasia. Annotations were performed blinded to molecular status. When feasible, the pathologist also provided a region-level assessment of fusion status. Tiles were sampled by regions of 33 grids of tiles to preserve contextual architectural features and ensure representative coverage.

| Cohort | A |
| --- | --- |
| Nb of Slides with Annotation | 15 |
| Nb of Regions Annotated (3x3 tiles) | 82 |
| Nb of Tiles Annotated per Slide: median (IQR) | 18.0 (54.0) |
| Nb of Tiles **annotated** | 738 |
| Nb of Tiles **fusion-negative / fusion-positive** (*Annotation by pathologist*) | 198 / 243 |
| Nb of Tiles with **monomorphous ovoid nuclei** | 292 |
| Nb of Tiles with **endocrinoid network of thin capillaries** | 116 |
| Nb of Tiles with **nuclear palisading** | 76 |
| Nb of Tiles with **presence of thin cytoplasmic cell processes between tumor nuclei and vessels / pseudorosette** | 42 |
| Nb of Tiles with **calcification** | 27 |
| Nb of Tiles with **desmoplasia** | 0 |

**Supplementary Figure 4.**
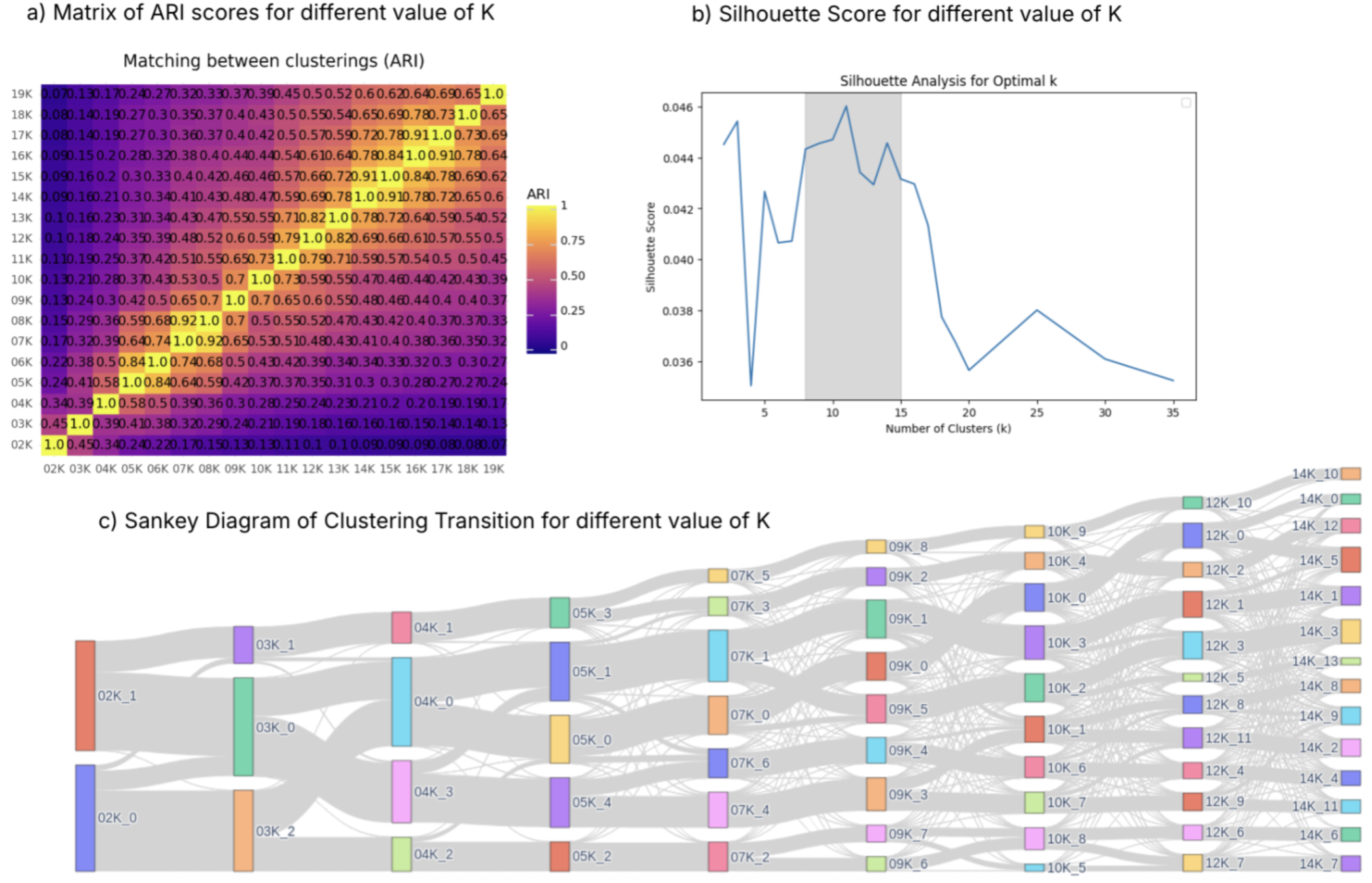
Selection of the number of clusters K leading to the number of tissue regions identified. **(a)** ARI between every pair of clustering obtained with a different number of clusters on the training cohort. **(b)** Silhouette scores for different numbers of tissue clusters on the training dataset. Interval of containing near maximal Silhouette score is marker in grey **(c)** Sankey diagram of the clustering with increasing number of clusters on the training dataset.

**Supplementary Figure 5.**
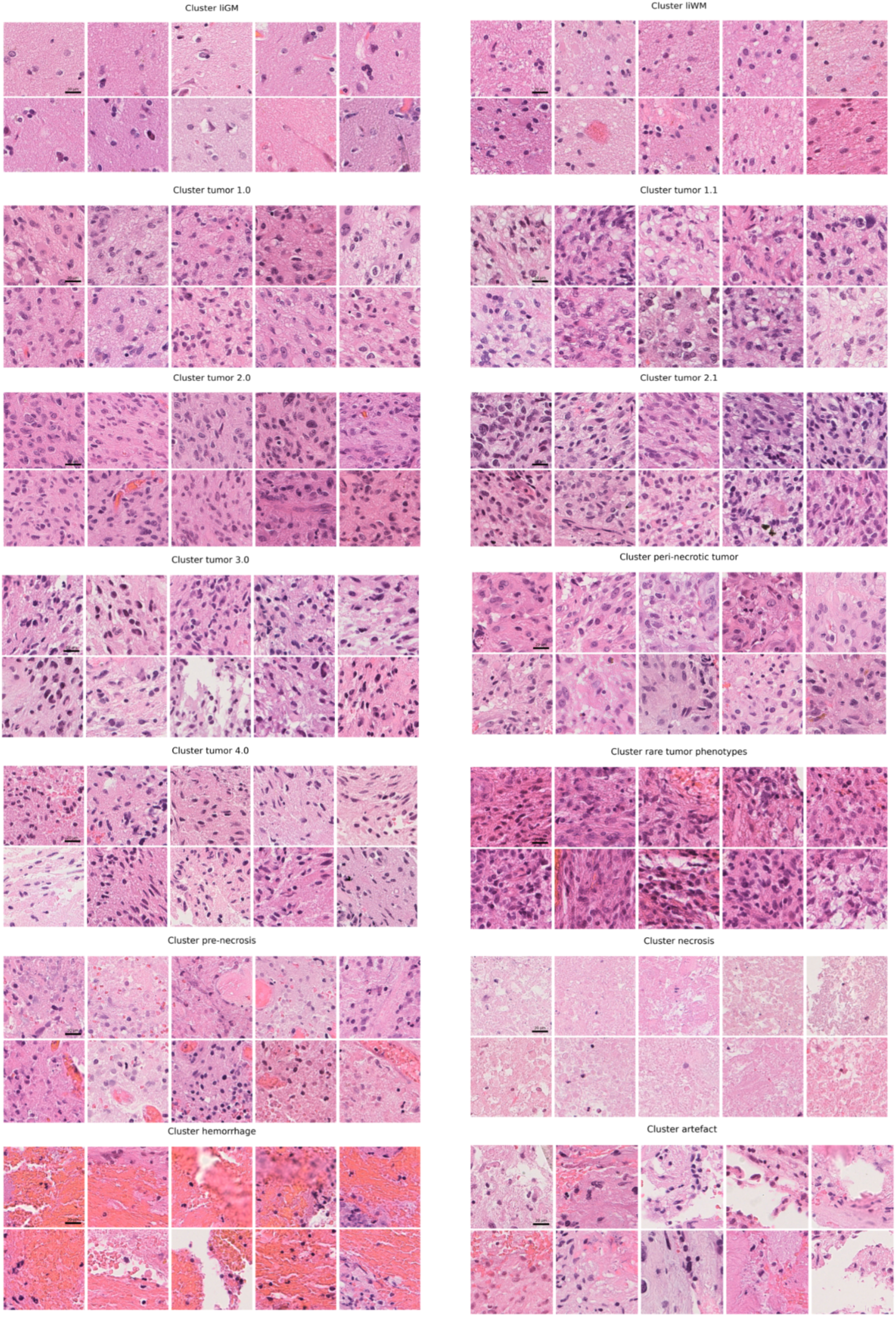
Illustration of the most representative tiles for each tissue region. Illustration of the 10 most representative tiles for each tissue subtypes. Most representative tiles correspond to the tiles with the smallest euclidean distance to the clusters centroids in the feature space. For each cluster, the 10 closest tiles are represented and sampled from 10 different slides. Table 2 provides descriptions of each cluster.

**Supplementary Figure 6.**
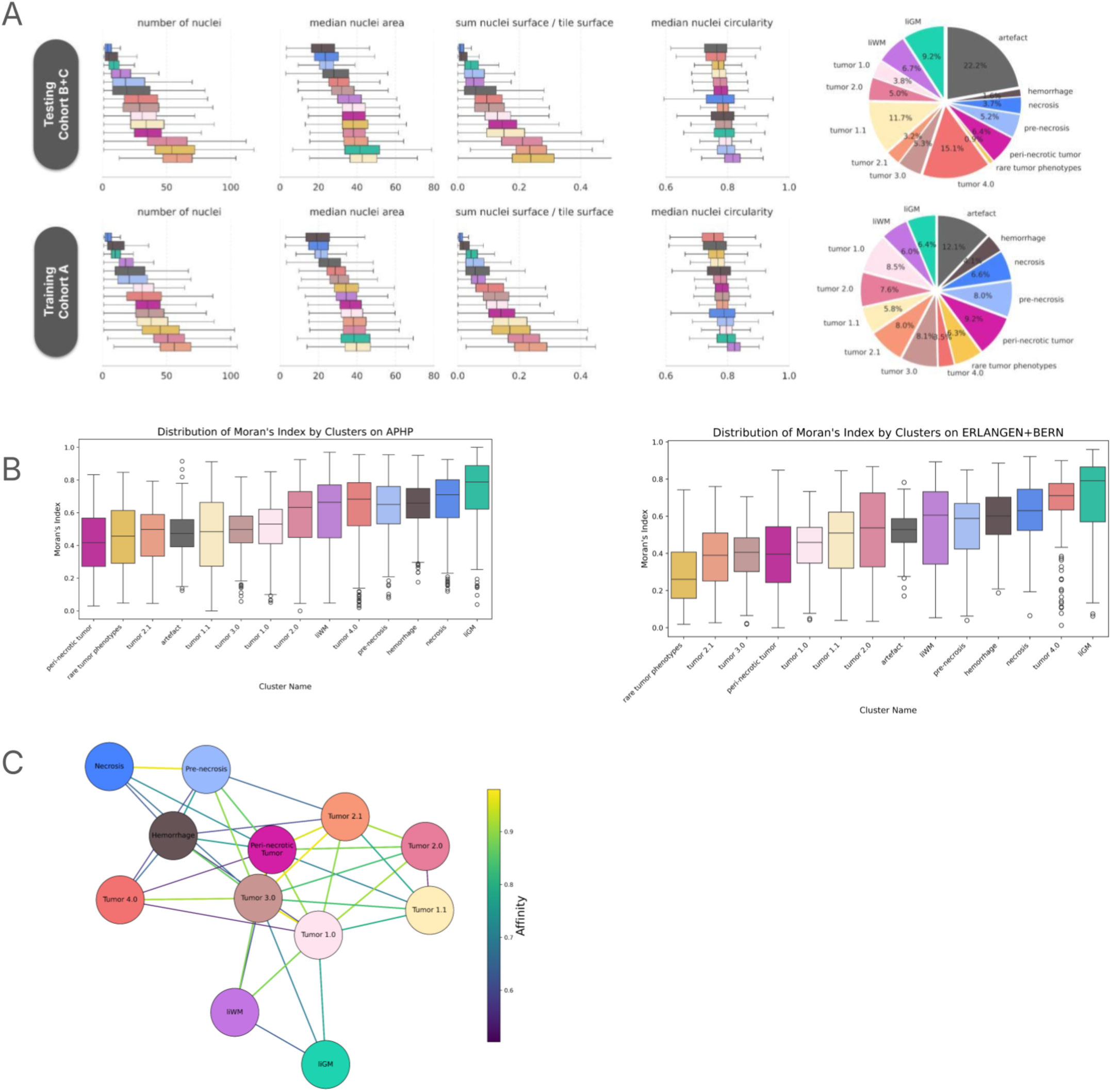
Characterization of the tissue regions observed in H&E slides across cohorts. **(a)** Number of nuclei, median nuclei area, the ratio of the sum of nuclei surface and the tile surface, and the median nuclei eccentricity for each tile of the H&E slides in the training (cohort A) and validation datasets (cohort B and cohort C). The box shows the quartile of the distributions while the whiskers extend to show the rest of the distribution except the outliers. Outliers were defined as points below the Q1 - 1.5 IQR or above Q3 + 1.5 IQR. **(b)** Distribution of Moran’s I per tissue regions computed on the training dataset. High Moran’s I indicates a more spatially compact tissue subtype. Tumoral tissue regions have lower Moran’s index than other regions, indicating that they are less spatially consistent and more intertwined with each other than the other. Yet all Moran’s I are positive. **(c)** Graph interpretation of tissue regions organization. Graph computed using the force-directed Fruchterman-Reingold algorithm for the tissue regions affinity matrix defined as the weighted average across the slides of the training cohort and the validation cohort of the Hausdorff distance at 5% between tissue regions within each slide. Low affinity values below 0.5 were clipped to 0 and were not displayed.

**Supplementary Figure 7.**
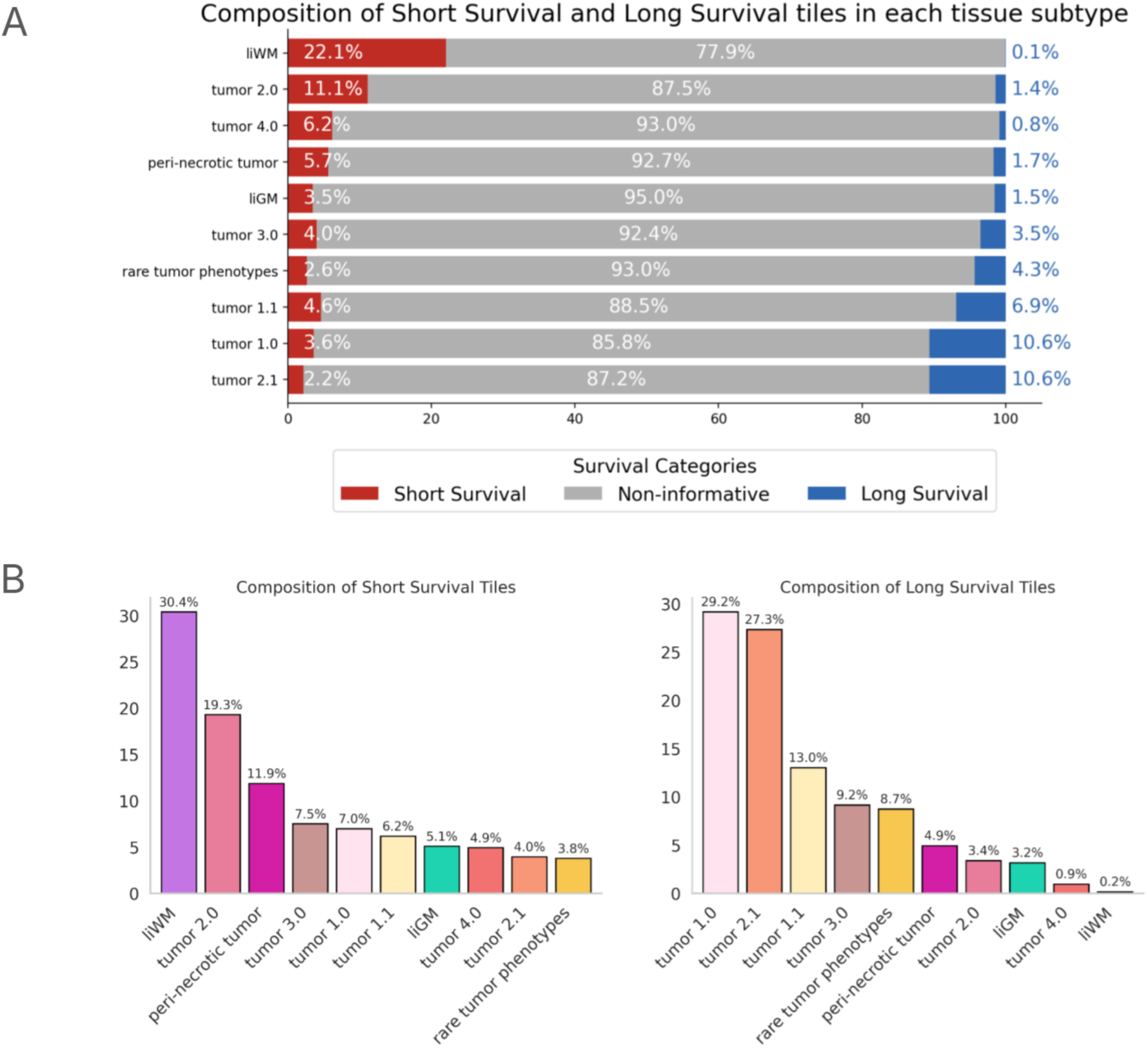
Composition of tiles associated with short survival, long survival, and non-informative categories in training dataset (Cohort A) **(a)** For each tissue subtype, proportion of tiles associated with short survival, long survival, and non-informative categories in Cohort A. **(b)** Distribution of tissue subtypes among Long Survival and Short Survival tiles in Cohort A

**Supplementary Figure 8:**
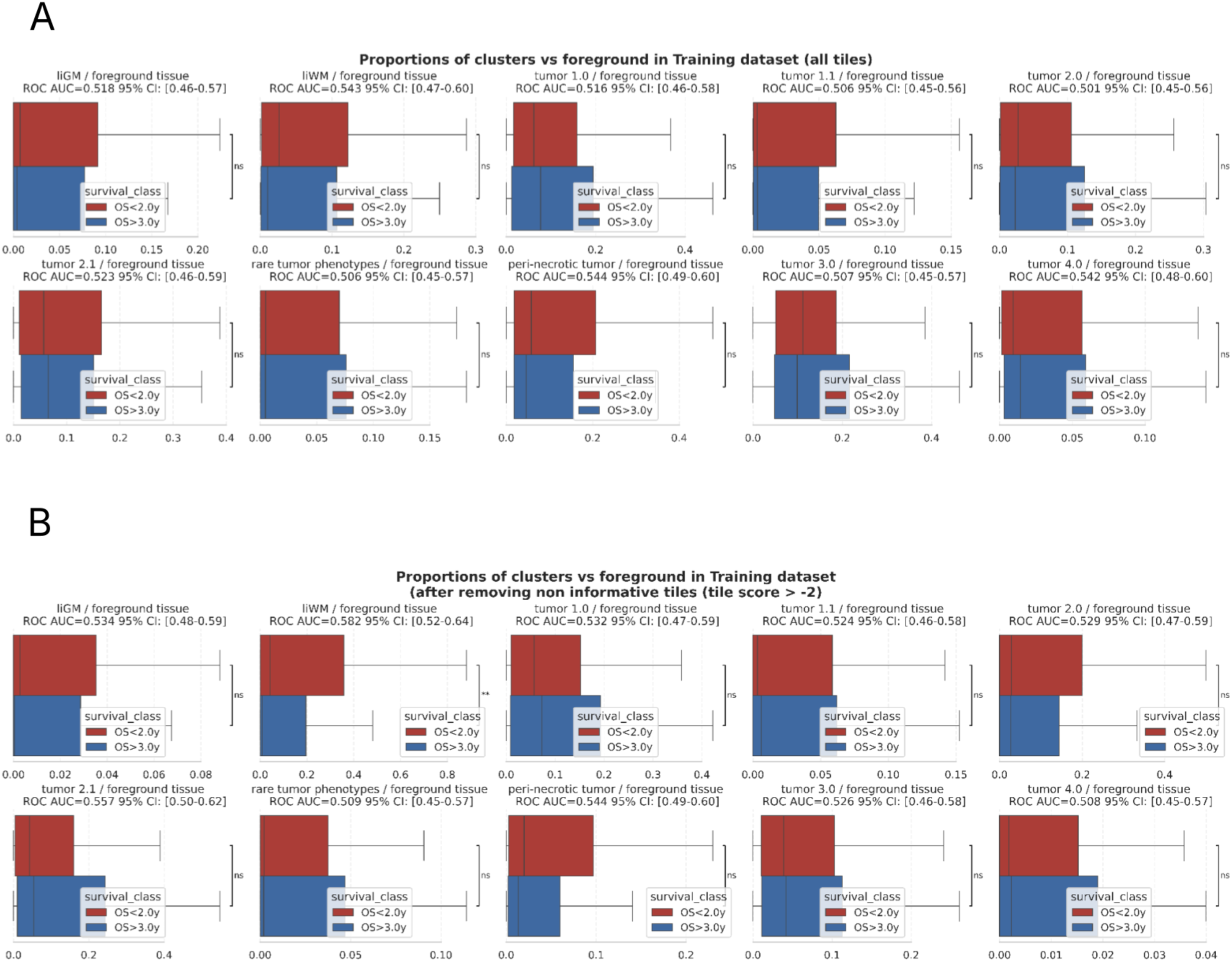
Tissue regions proportions alone do not predict survival. **(a)** Distribution of cluster proportions relative to foreground tissue across all tiles for each slide. Foreground tissue includes all subtypes except necrosis, pre-necrosis, hemorrhage, and artefact. **(b)** Same as **(a)**, but excluding non-informative tiles (tile score < –2). Threshold optimization is detailed in Supplementary Figure 15. For each cluster, ROC AUC values quantify the association between tissue proportion and survival class. Confidence intervals are estimated via 1,000 bootstrap iterations. Statistical significance is assessed using a two-sided Mann–Whitney U test. P values are indicated as follows: *** < 0.001; ** < 0.01; * < 0.05; ns, not significant.

**Supplementary Figure 9.**
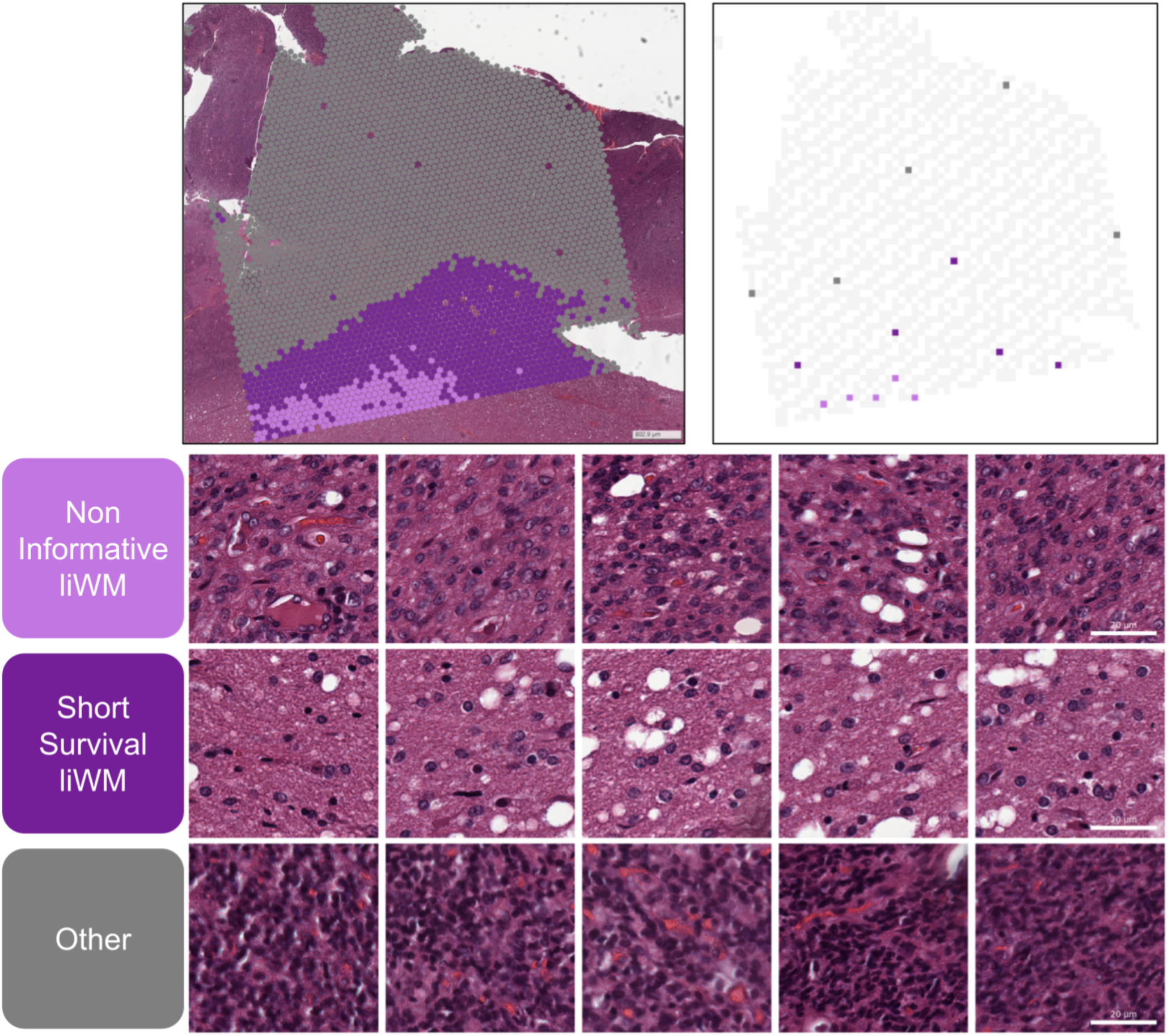
Illustration of the liWM short survival biomarke. Illustration of high resolution images of the liWM short survival biomarker on a VISIUM sample. The top right panel indicates tiles sampling location.

**Supplementary Figure 10.**
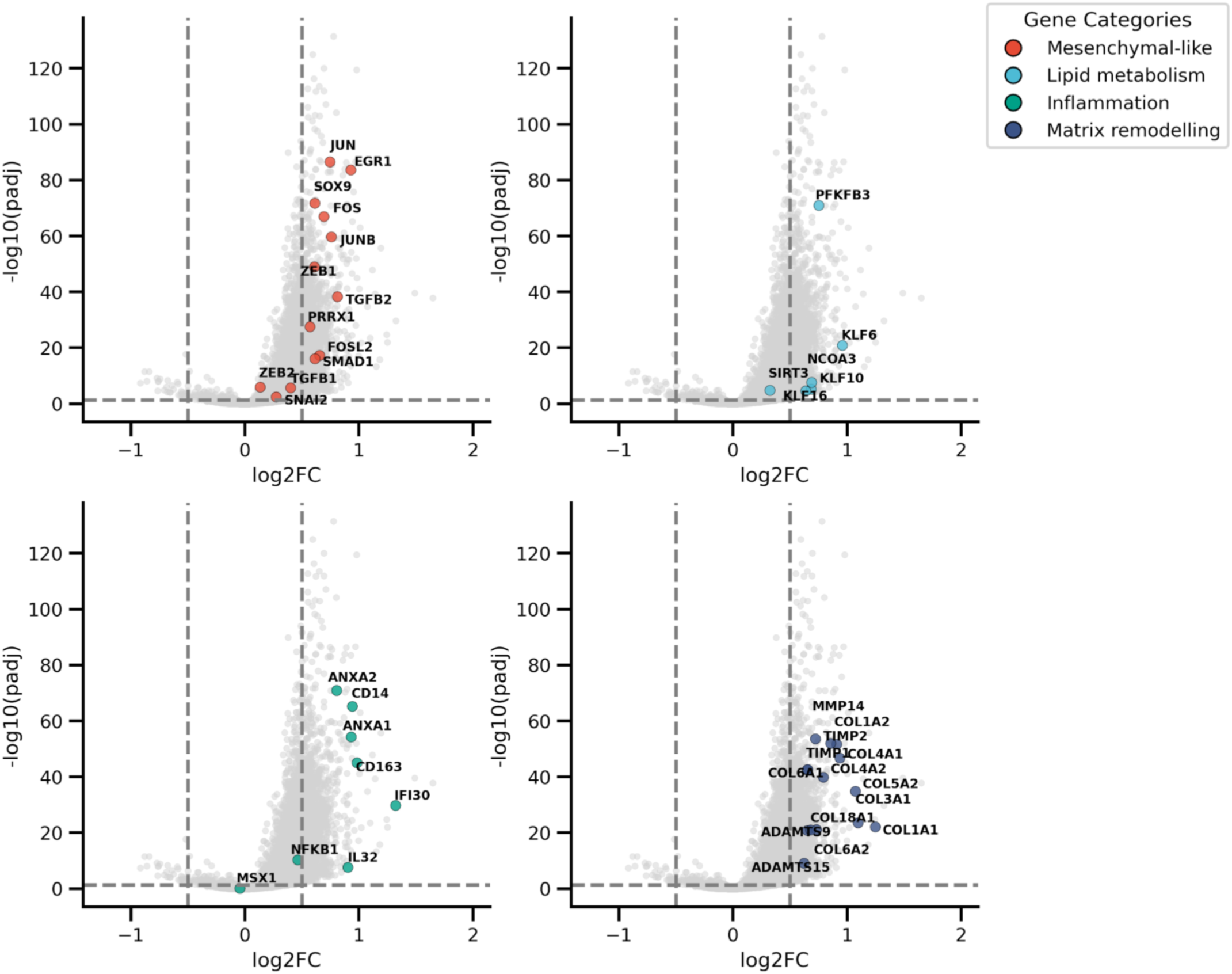
Volcano plot with highlighted genes. The list of genes is available in supplementary Table K.

**Supplementary Figure 11.**
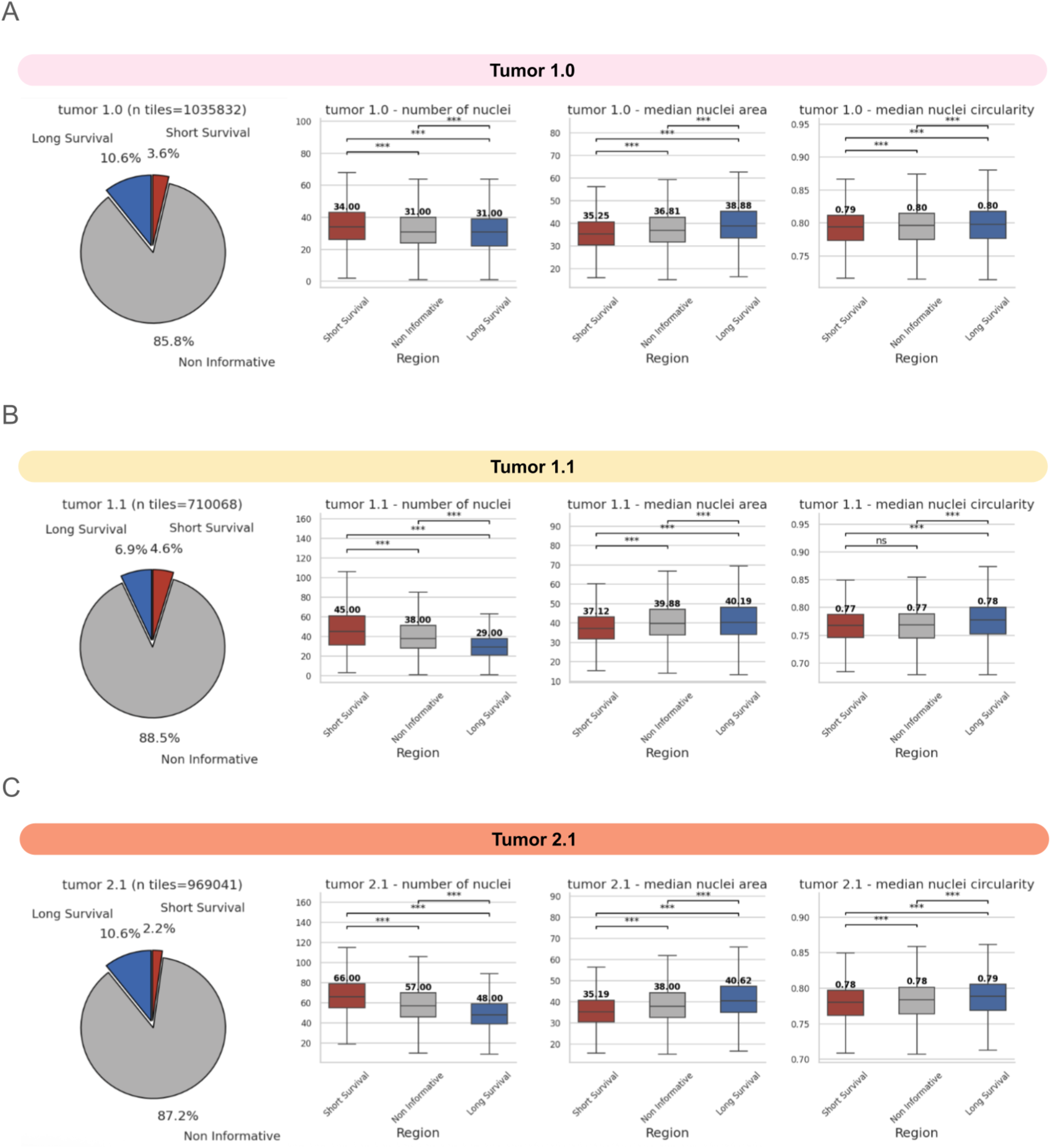
Morphological differences between tiles of short survival, long survival, and non-informative regions in clusters enriched in long survival regions: tumor 1.0 (a), tumor 1.1 (b), and tumor 2.1 (c). Cell-level features were computed on the training dataset (cohort A).

**Supplementary Figure 12.**
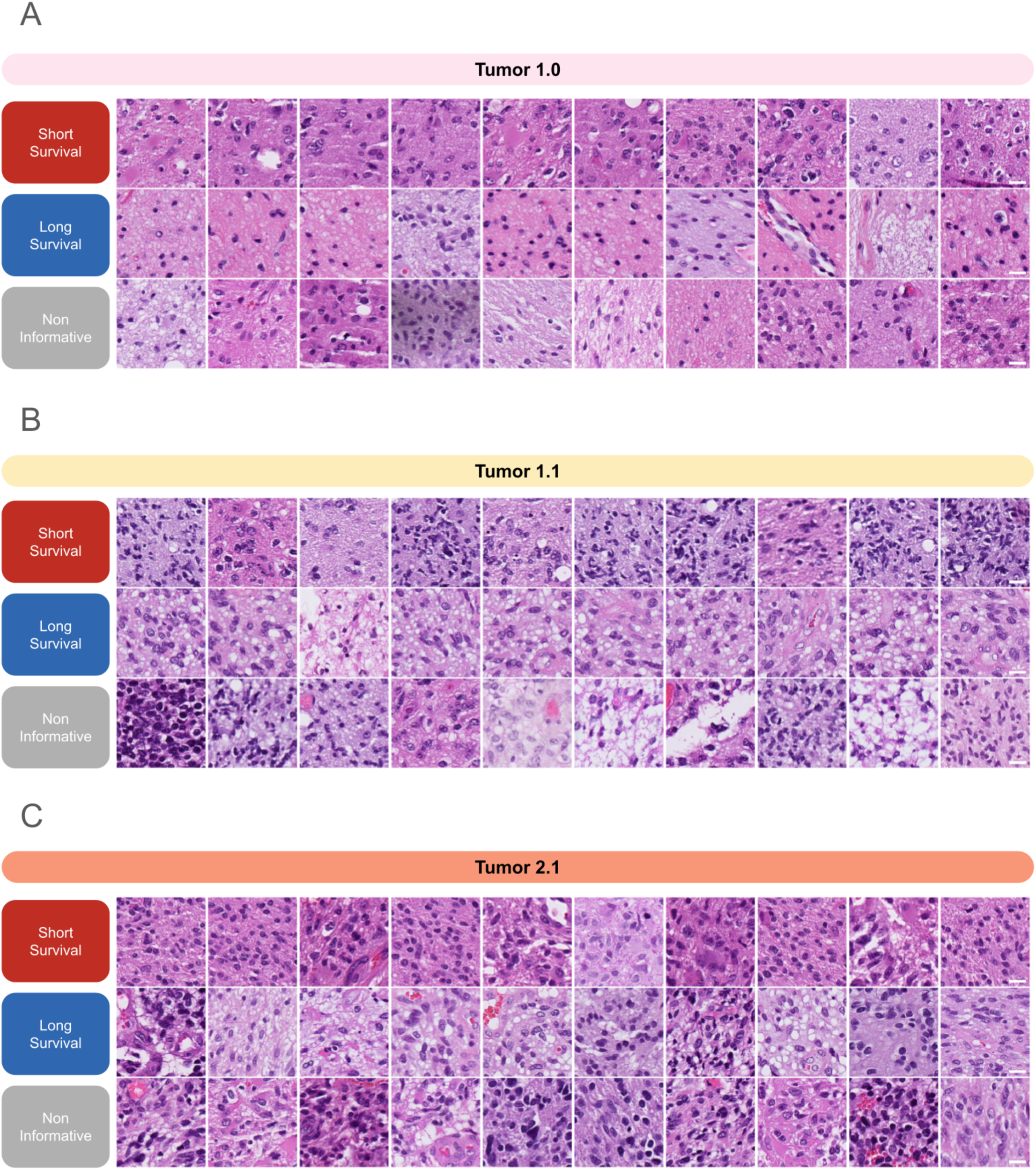
Illustration of tiles of Short survival, Long survival, and Non-informative regions within the tumor tissue regions in the validation dataset (cohort B & C). Tiles are selected randomly within each sub-category (short-survival, long-survival, non-informative) after pooling all the tiles from cohorts B & C.

Supplementary Table H. Top genes of the 41 cell types analysed in Spatial Transcriptomics.

***Due* to *its size, Supplementary Table H is provided in the Supplementary Data and cannot be rendered in the main manuscript*.**

Supplementary Table I. Pathways enriched in short survival liWM vs non informative liWM.

***Due* to *its size, Supplementary Table*** *I* ***is provided in the Supplementary Data and cannot be rendered in the main manuscript*.**

Supplementary Table J. Genes enriched in short survival liWM vs non informative liWM.

***Due* to *its size, Supplementary Table* J *is provided in the Supplementary Data and cannot be rendered in the main manuscript*.**

Supplementary Table K. Volcano plots (each signature highlighted in an individual volcano plot)

***Due* to *its size, Supplementary Table J is provided in the Supplementary Data and cannot be rendered in the main manuscript*.**

Supplementary Table L. Pathways enriched in long vs. short-survival.

***Due* to *its size, Supplementary Table J is provided in the Supplementary Data and cannot be rendered in the main manuscript*.**

**Supplementary Table M.** Blood vessels manual annotation: description per cohort. Number of tiles annotated by an expert pathologist FB for presence of blood vessels and the five blood vessel subtypes: hyperplasic endothelium capillary (HEC), thin endothelium capillary (TEC), microvascular proliferation (MVP), hyperplasic endothelium with wide lumen (HEWL), and thin endothelium with wide lumen (TEWL).

| Cohort | Nb of Slides with Annotation | Nb of Tiles Annotated per Slide: median (IQR) | Nb of Tiles with/without vessels | Nb of Tiles with HEC | Nb of Tiles with TEC | Nb of Tiles with MVP | Nb of Tiles with HEWL | Nb of Tiles with TEWL |
| --- | --- | --- | --- | --- | --- | --- | --- | --- |
| A | 80 | 6 (5.25) | 816/928 | 280 | 327 | 83 | 22 | 76 |
| B | 15 | 13.5 (10.25) | 245/280 | 106 | 80 | 3 | 19 | 37 |

**Supplementary Table N.**
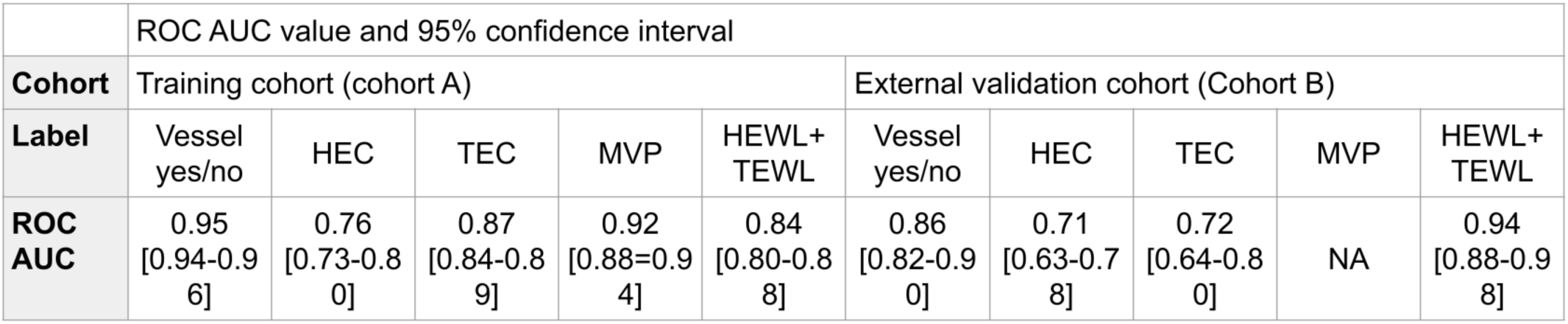
Performance of models for blood vessels detection and classification. Performance of our H&E models (ensemble of logistic regression models) for predicting the presence of vessel (model 1) of the vessel subtypes (model 2). For model 2, HEWL and TEWL were merged due to the low number of annotations available for training. ROC AUC are computed with a one-versus-all approach. ROC AUC was not reported for the MVP on the external validation cohort due to the low number of MVP that was annotated in this cohort (n=3).

**Supplementary Figure 13.**
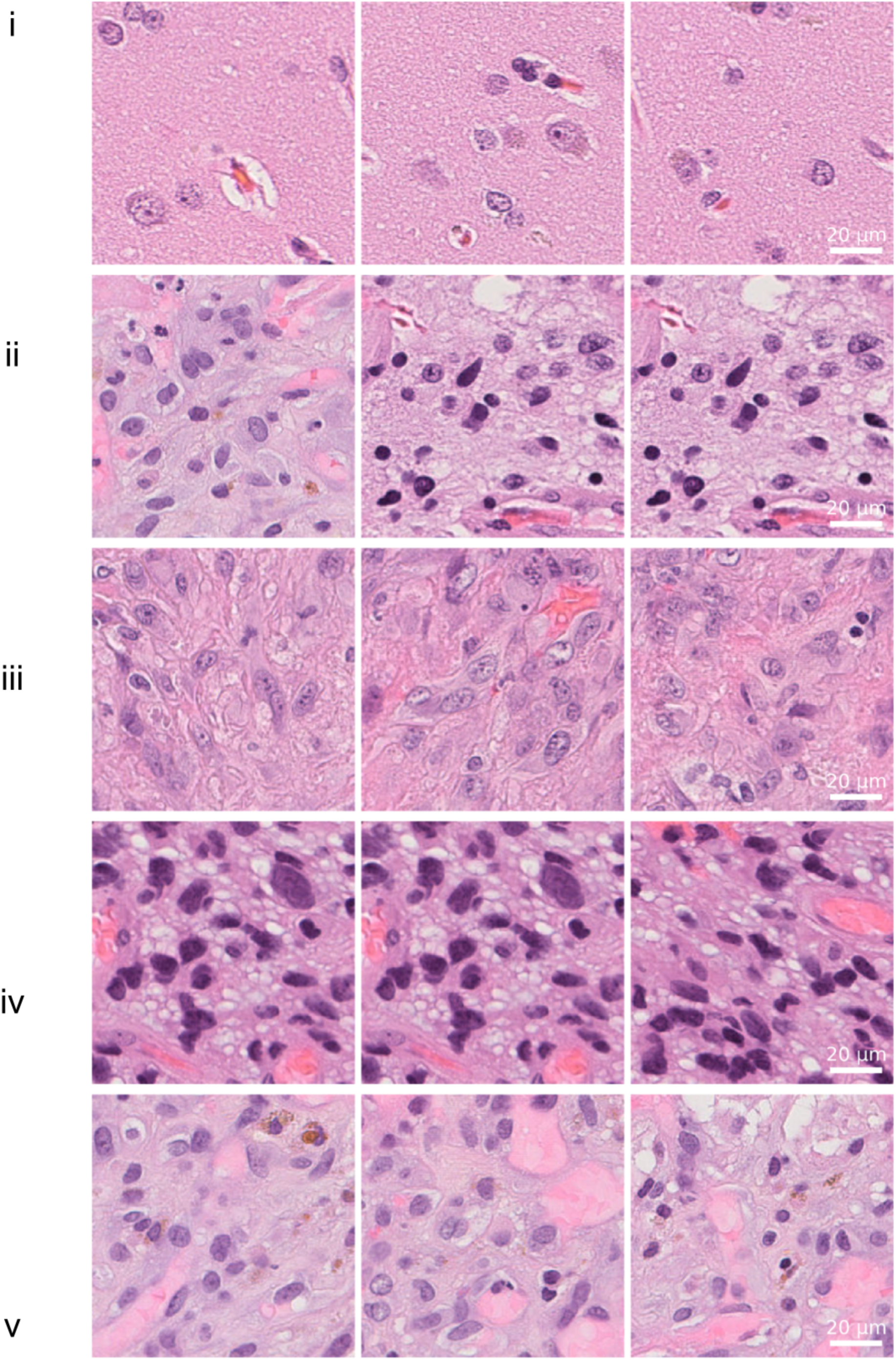
Examples of regions illustrating the different annotated vessel patterns. (i) Thin Endothelium Capillary (TEC) - one red blood cell (ii) Hyperplasic Endothelium Capillary (HEC) - one red blood cell (iii) Microvascular Proliferation (MVP) (iv) Thin Endothelium Wide Lumen (TEWL) - multiple red blood cells (v) Hyperplasic Endothelium Wide Lumen (HEWL) - multiple red blood cells

**Supplementary Figure 14.**
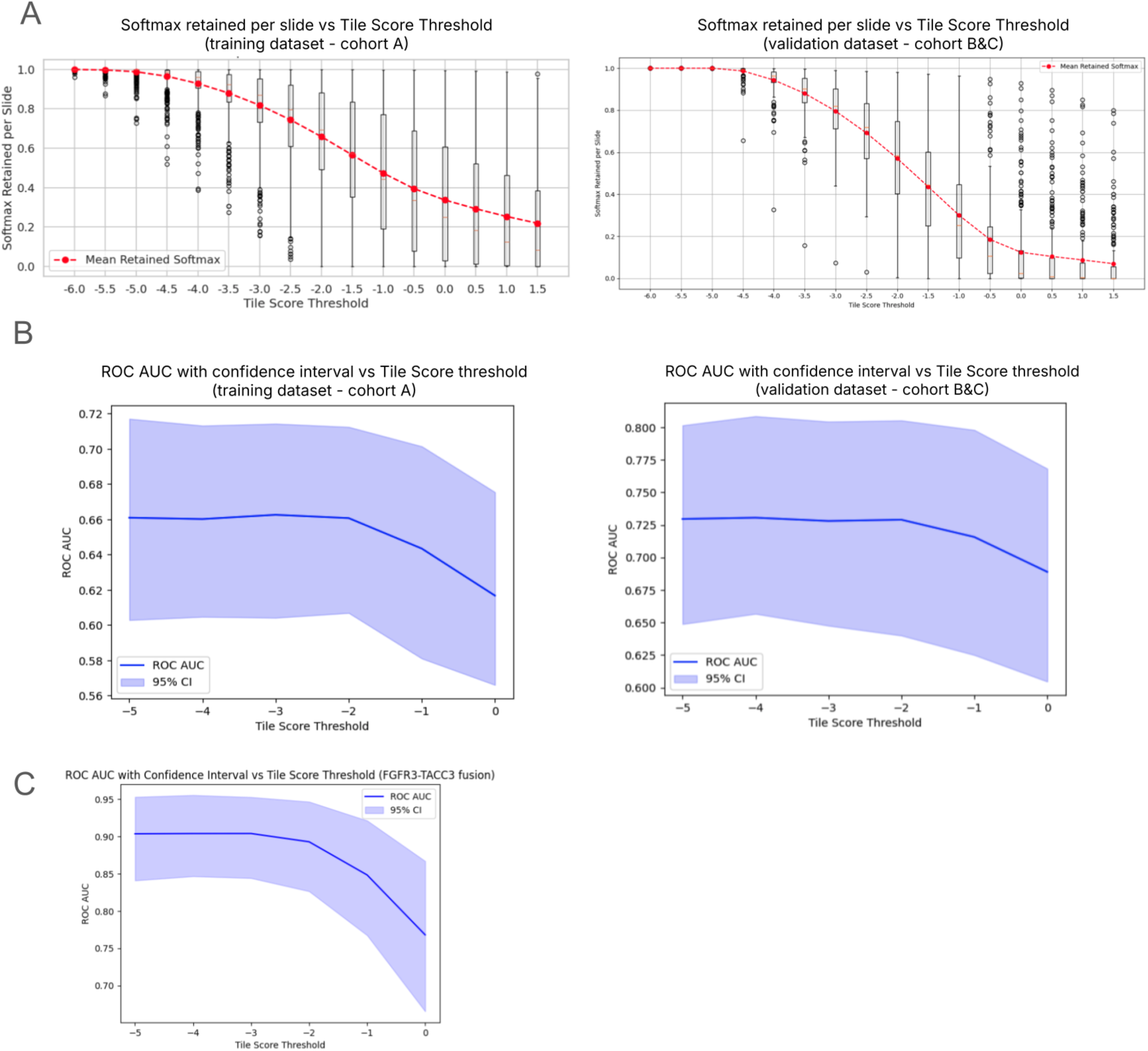
Selection of the tile score threshold. **(a)** Percentage of softmax Tile Scores per Slide retained for different Tile Score thresholds in Cohort A (left) and Cohort B and C (right). **(b)** Evolution of ROC AUC of the H&E tile predictions model with different values of tile score threshold on Cohort A (left) and Cohort B and C (right) for the OS classification task **(c)** Evolution of ROC AUC of the H&E tile predictions model with different values of tile score threshold on Cohort A (left) for the FGFR3-TACC3 fusion classification task.

